# An evolutionary-conserved redox regulatory mechanism in human Ser/Thr protein kinases

**DOI:** 10.1101/571844

**Authors:** Dominic P. Byrne, Safal Shrestha, Natarajan Kannan, Patrick A. Eyers

## Abstract

Reactive oxygen species (ROS) are products of oxygen metabolism, but are also recognized as endogenous physiological mediators of cellular signaling. Eukaryotic protein kinase (ePK) regulation occurs through reversible phosphorylation events in the flexible activation segment. In this study, we demonstrate that the catalytic phosphotransferase output from the mitotic Ser/Thr kinase Aurora A is also controlled by cysteine (Cys) oxidation. Reversible regulation occurs by direct modification of a conserved residue (Cys 290), which lies adjacent to Thr 288, the activating site of phosphorylation. Strikingly, redox modulation of the Cys 290-equivalent in other ePKs is predicted to be an underappreciated regulatory mechanism, since ~100 human Ser/Thr kinases possess a Cys at this position in the conserved activation loop. Using real-time enzyme assays, we confirm that the presence of the equivalent Cys residue is prognostic for redox-sensitivity amongst a cohort of human CAMK, AGC and AGC-like kinases, including AKT, AMPK, CAMK1, MAPKAP-K2/3 and SIK1-3. Our findings demonstrate that dominant Cys-based redox-switching in the activation segment represents an evolutionary-conserved mode of regulation for a significant subset of the human kinome. This finding has important implications for understanding physiological and pathological signaling responses to ROS, and emphasises the importance of multivalent activation segment regulation in ePKs.

**ONE-SENTENCE SUMMARY:** The catalytic activity of Ser/Thr kinases is regulated through a conserved Cys-based redox mechanism.

## INTRODUCTION

Protein phosphorylation on Ser/Thr and Tyr residues controls all cellular aspects of eukaryotic life. In order to regulate the flow of signaling information, the enzymes that catalyse the addition and removal of phosphate groups are themselves subject to reversible regulation. In the case of Ser, Thr and Tyr kinases, this often involves phosphorylation-dependent mechanisms in which conserved residues are cyclically phosphorylated and dephosphorylated to control catalysis. A well-known example is the reversible phosphorylation of Ser/Thr and Tyr residues in the conformationally-flexible ‘activation segment’, which can either be liberated or fold-back onto the kinase to inhibit substrate phosphorylation (Nolen et al., 2004). The activation segment, also known as the T-loop, is located between the DFG and APE motifs (Johnson et al., 1996), two highly-characteristic regions found in most canonical ePKs (Kannan and Neuwald, 2005). In the case of pTyr regulation, redox control of Tyr phosphatases is well-documented (Salmeen et al., 2003, van Montfort et al., 2003, Tonks, 2005), providing an extra layer of regulation in addition to reversible phosphoregulation of individual Tyr kinases. For example, the oxidation of an acidic Cys residue in the catalytic motif of the tyrosine phosphatase PTP1B to a cyclic sulfenylamide inhibits catalysis, which can be reversed by reducing agents such as DTT that regenerate a Cys sulfhydryl (Salmeen et al., 2003, van Montfort et al., 2003). In addition, regulation of Ser/Thr phosphatase signalling components has been proposed to be important in oxidation-regulated cell cycle transitions (Lim et al., 2015, Savitsky and Finkel, 2002).

Reactive oxygen species (ROS), the collective term for reactive oxygen-derived radicals that include superoxide and peroxide, were originally regarded as side-products of oxygen metabolism, but are now recognized as important endogenous regulators of cellular signaling across the cell cycle (Paulsen and Carroll, 2010, Patterson et al., 2019, Seo and Carroll, 2009). The sulfur atom of Cys residues, the predominant intracellular redox-signaling molecule, can exist in a variety of interconvertable oxidation states in redox active signaling proteins (fig. S1)(Rhee et al., 2005). In cells, oxidation of a reactive cysteine thiolate anion (Cys-S^−^) results in the formation of the transient sulfenic acid species (Cys-SOH), which can either undergo further (irreversible) oxidation to stable sulfinic (Cys-SO_2_H) or sulfonic (Cys-SO3H) acid, or be recycled by antioxidants constituting the oxidative stress response (Gupta and Carroll, 2014). An important example is that of a reversible catalytic sulfenamide intermediate (fig. S1), which is generated in oxidized versions of the tyrosine phosphatase PTP1B (Salmeen et al., 2003, van Montfort et al., 2003) and reduced by cellular agents such as glutathione (Forman et al., 2017).

In both prokaryotes and eukaryotes the reduced glutathione (GSH) pool acts as a buffer of the cellular environment, with physiological concentrations ranging from ~1 to 10 mM (Grek et al., 2013, Anselmo and Cobb, 2004, Schafer and Buettner, 2001, Owen and Butterfield, 2010, Maher, 2005). In the cytosol, glutathione exists as oxidized (GSSG) and reduced (GSH) species, and the GSSG/GSH ratio changes as a function of redox stress (Owen and Butterfield, 2010). Reversible modification of protein cysteine thiol groups through disulfide bond formation with GSH is widely considered to be a defence mechanism, which protects proteins against proteotoxic stress from over-oxidation (Dalle-Donne et al., 2007).

For ePKs, several examples of redox-regulatory mechanisms have been reported, although the lack of a catalytic Cys in the kinase active site means that these mechanisms are centred on other regions of the kinase domain, most notably modification of conserved Cys residues in the Gly-rich loop. Examples of redox-sensitive Tyr kinases include ABL, SRC, EGFR and FGFR (Truong and Carroll, 2012, Heppner et al., 2018, Kemble and Sun, 2009, Giannoni et al., 2005). Related work, submitted back-to-back with this manuscript, demonstrates that the evolutionary-related prokaryotic and eukaryotic Fructosamine 3 kinases (FN3Ks), which lack the conventional activation segment found in ePKs, also employ reversible Cys regulatory mechanisms to control catalytic output (Shrestha, 2019). Like the Tyr kinases, FN3K’s possess conserved redox-active Cys residues in the Gly-rich loop that function as oxidizable switches to co-ordinate reversible dimerization and control enzymatic (in)activation.

This discovery of chemically-accessible (and redox-sensitive) Cys residues in protein kinases has created new opportunities for the design of chemical reagents and covalent clinical compounds to target these residues with specificity (Butterworth et al., 2017, Zhao et al., 2017, Liu et al., 2013, Byrne et al., 2017, Chaikuad et al., 2018). Indeed, several examples of Ser/Thr kinase regulation by the modification of redox-active Cys residues exist, although no overarching evolutionary-based mechanism has been proposed. Examples of redox-regulated proteins containing kinase domains include ASK1, MEKK1, MELK, PKA, PKG, ERK, JNK and p38 MAPK-family members (Cuello and Eaton, 2019, Cross and Templeton, 2004, Corcoran and Cotter, 2013a, Burgoyne et al., 2007b, Nadeau et al., 2009, Humphries et al., 2007b, Shao et al., 2014, Cao et al., 2013b, Murata et al., 2003b) although, the mechanism of redox modulation of PKG Cys residues, both inside and outside of the catalytic domain, remains controversial (Kalyanaraman et al., 2017, Burgoyne et al., 2013, Sheehe et al., 2018, Cuello and Eaton, 2018, Prysyazhna et al., 2012, Burgoyne et al., 2007a). Indeed, without a common set of biochemical reagents and reliable real-time assay conditions, it is currently challenging to define common themes for redox-based regulation amongst protein kinases.

Aurora A is an oncogenic Ser/Thr protein kinase (Bischoff et al., 1998), which is subject to multi-level reversible regulation in human cells including phosphorylation/dephosphorylation in the activation segment and allosteric control by accessory factors such as TPX2, TACC3 and protein phosphatases (Eyers et al., 2003, Bayliss et al., 2003, Eyers and Maller, 2004, Burgess et al., 2018). Aurora A controls the G2/M transition (Macurek et al., 2008) and is also required for centrosome separation and mitotic spindle assembly (Carmena and Earnshaw, 2003, Hegarat et al., 2011). Aurora A-dependent signaling is also implicated in mitochondrial dynamics and metabolism, which are closely associated with the production of ROS (Bertolin et al., 2018). Recent work has identified the structural and biochemical basis of a novel mode of Cys-based inhibition in Aurora A (Tsuchiya, 2018), which occurs through an interaction between Cys 290 in the activation segment and Coenzyme A (CoA). Aurora A activity is reversibly inhibited by CoAlation on Cys 290, which also occurs in human cells exposed to oxidative stresses (Tsuchiya, 2018). However, these studies did not reveal whether Aurora A catalytic activity is regulated by classical redox agents in human cells.

In this study, we report that the catalytic activity of Aurora A, an essential cell cycle-regulated Ser/Thr kinase, is controlled by reversible Cys oxidation. DTT-reversible Aurora A oxidation by a variety of agents occurs through modification of Cys 290, which lies adjacent to Thr 288, the established activating ‘T-loop’ site of phosphorylation. This Cys residue is conserved in Aurora kinases throughout eukaryotic evolution, and redox modulation of the Cys 290-equivalent in eukaryotic kinases appears to be a dominant, and evolutionary-conserved, mechanism since ~10% of 285,479 protein kinase-related sequences in the non-redundant (NR) sequence database contains this conserved Cys residue. These include well-known kinases such as PKA, PLK1 and PLK4, which we confirm are all regulated by a reversible Cys-based redox mechanism. We also establish that the presence of a Cys residue is prognostic for redox-sensitivity amongst human CAMK and AGC-like kinases, but not Tyr kinases, which lack Cys in the activation segment. Important examples of redox-regulated Ser/Thr kinases include AKT, SIK, AMPK, MAPKAP-K2/3 and CAMK1 proteins. Overall, our study demonstrates that redox modification of the activation segment represents a conserved extra layer of regulation for subsets of eukaryotic protein kinases, including a significant proportion of the human kinome. These findings have important implications for understanding physiological and pathological responses to ROS in cells, and help rationalise numerous lines of experimental evidence demonstrating that reducing agents are required for the catalytic activity of many, but not all, Ser/Thr kinases.

## RESULTS

### Redox regulation of Aurora A

Human Aurora A was purified to homogeneity in the presence or absence of DTT, and immunoblotting with a phosphospecific antibody confirmed similar levels of pThr 288 in both enzyme preparations, as expected for active, autophosphorylated, Aurora A (fig. S2A). Regardless of the source of the enzyme, Aurora A activity was inhibited by H_2_O_2_ in a concentration dependent manner, with ~70 % inhibition observed at 100 μM H_2_O_2_ (Fig. 1A). Diamide, which also oxidizes exposed cysteine residues in proteins (Kosower and Kosower, 1995, Humphries et al., 2002), also inhibited Aurora A activity (Fig. 1A). In contrast, exposure of Aurora A to an excess of DTT enhanced catalytic activity, regardless of the source of enzyme (compare Fig. 1A and fig. S2B). To further evaluate the redox response of Aurora A, we employed a physiological Aurora A substrate, TACC3, and confirmed that phosphorylation at Ser 558 was inhibited in a dose-dependent manner by H_2_O_2_, with low mM concentrations of peroxide blocking activity as efficiently as the Aurora A inhibitor MLN8237 (Fig. 1B). In contrast, DTT markedly increased phosphotransferase activity towards the TACC3 substrate (Fig. 1B). Next, we sought to establish whether oxidation by H_2_O_2_ resulted in irreversible inhibition of the kinase. Phosphorylation of the peptide substrate by Aurora A was monitored in real time in the presence or absence of H_2_O_2_ prior to the addition of DTT. After 40 mins, only ~ 15 % of the peptide substrate had been phosphorylated by Aurora A when H_2_O_2_ was included in the reaction, compared to ~80% in its absence (Fig. 1C). However, supplementing the reaction with DTT restored oxidized Aurora A activity, resulting in an immediate increase in the rate of substrate phosphorylation compared to an experiment maintained in an oxidized (inhibited) state (Fig. 1C). The activity of Aurora A purified in the presence of DTT could also be rescued in real-time by the addition of DTT following inhibition by H_2_O_2_ (fig. S2C). These findings indicate that inhibition of Aurora A by oxidative modification is reversible and suggest that Aurora A catalytic activity can be regulated by its redox state.

**Figure 1.**
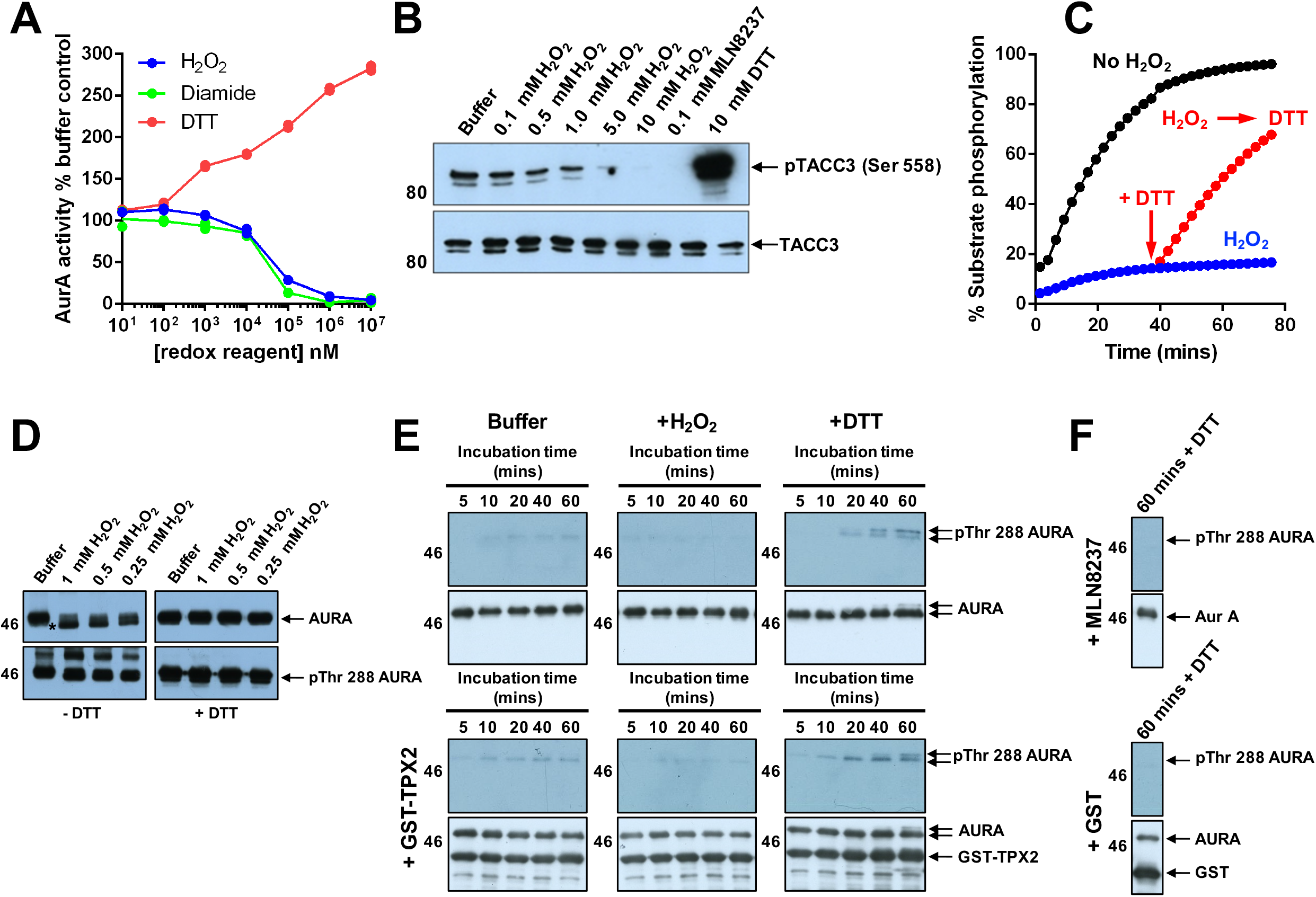
Redox-dependent regulation of Aurora A activity *in vitro*. **(A)** Dose response curves for the reducing reagent DTT (red) and the oxidizing agents H_2_O_2_ (blue) and diamide (green) with 6 nM recombinant Aurora A in the presence of 1 mM ATP. Aurora A activity was assessed by monitoring phosphorylation of a fluorescent peptide substrate, and normalized to controls after 30 min assay. **(B)** Immunoblot of an in vitro kinase assay using recombinant GST-TACC3 as a substrate for Aurora A. GST-TACC3 (1 μg) was incubated at 20 °C for 10 mins with 3 ng Aurora A with 0.5 mM ATP and 5 mM MgCl_2_. Aurora A-dependent phosphorylation of TACC3 (pSer558, top panel) was detected in the presence of the indicated concentrations of H_2_O_2_, 10 mM DTT or 0.1 mM of MLN8237. Reactions were terminated by the addition of SDS loading buffer. Equal loading of TACC3 substrate was confirmed with anti-TACC3 antibody (bottom panel). **(C)** Oxidative-inhibition of Aurora A is reversible. Aurora A (12.5 nM) activity was monitored in real time in the presence (blue) or absence (black) of 1 mM H_2_O_2_. After 40 mins, reactions were supplemented (where indicated) with 2 mM DTT (red). Aurora A dependent phosphorylation of the fluorescent peptide substrate was monitored using assay conditions described in (A). **(D)** Immunoblot demonstrating reversible increase in the electrophoretic mobility of Aurora A, presumably due to oxidation by H_2_O_2_. Aurora A (0.5 μg) was incubated with the indicated concentrations of H_2_O_2_ for 10 mins at 20°C and analysed after non-reducing or reducing SDS-PAGE. Asterisk denotes the reversibly oxidized species. Total Aurora A (upper panel) and pThr 288 Aurora A (lower panel) blots are shown. (E, F) Immunoblot demonstrating redox-dependent Aurora A autophosphorylation at Thr 288. Dephosphorylated Aurora A (1 μg) was produced by co-expression with lambda phosphatase in *E.coli* and then incubated with 1 mM ATP and 10 mM MgCl2 for the indicated time periods under reducing (+ 1 mM DTT) or oxidizing (+ 1 mM H_2_O_2_) conditions in the presence and absence of TPX2 or MLN8237. Reactions were terminated by the addition of SDS loading buffer.

To rule out ‘non-specific’ kinase inactivation as a result of oxidative protein unfolding, we performed thermal stability measurements for Aurora A incubated in the presence of H_2_O_2_. The unfolding profile obtained for Aurora A was unaltered by inclusion of H_2_O_2_, DTT or reduced glutathione (GSH), with T_m_ values of ~40°C observed under test conditions (fig. S3A). Next, to evaluate the consequence of oxidation of Aurora A on its ability to bind ATP, we measured the rate of phosphate incorporation into our peptide substrate in the presence of different concentration of H_2_O_2_. Michaelis-Menten kinetic analysis revealed that H_2_O_2_ treatment significantly reduced the catalytic constant, *K_cat_*, of Aurora A, without affecting the affinity for ATP (inferred from K_M[ATP]_ values) (fig. S3B). Consistent with a lack of effect on the nucleotide-binding site, DSF analysis demonstrated essentially identical ΔT_m_ values induced by binding of ATP or the kinase inhibitor MLN8237 in the presence of H_2_O_2_, DTT or GSH (fig. S3C). Together, these data indicate that oxidation inhibits Aurora A ability to facilitate substrate phosphorylation without affecting the affinity for ATP or the thermal stability (unfolding) profile of the enzyme.

### Aurora A autoactivation is stimulated by DTT

One crucial prerequisite for Aurora A activation is autophosphorylation on Thr 288 in the activation loop (Littlepage et al., 2002, Walter et al., 2000a). To determine if Aurora A oxidation alters the extent of Thr 288 phosphorylation, it was incubated with increasing concentrations of H_2_O_2_ and analysed by western blotting under reducing or non-reducing SDS-PAGE conditions. Although the inclusion of H_2_O_2_ had no effect on the amount of phosphorylated Thr 288, oxidation altered the electrophoretic mobility in the absence of DTT resulting in a slightly faster migrating band (Fig. 1D, asterisk). Importantly, formation of this oxidation-dependent high mobility species of Aurora A was reversible, since it was abolished after reducing SDS-PAGE. A change in migration of oxidized proteins during SDS-PAGE is indicative of an oxidized Cys residue (Rudyk and Eaton, 2014, Sheehan, 2006). Although these findings clearly demonstrate the formation of a reversibly oxidized species of Aurora A, the depletion of kinase activity could not be attributed to a loss of Thr 288 phosphorylation. Next, we investigated the rate of autophosphorylation by Aurora A after co-expression with lambda phosphatase (λPP), which removes all activating phosphate from Thr 288. Accumulation of pThr 288 was assessed by immunoblotting following the addition of ATP and Mg^2^+ ions in the presence and absence of the known allosteric activator TPX2. Under standard assay conditions, Aurora A autophosphorylation was extremely slow, and only trace amounts of pThr 288 Aurora A were detected after 60 mins (Fig. 1E). In contrast, the inclusion of DTT (or TPX2) markedly increased the rate of Aurora A autophosphorylation, whereas no pThr 288 Aurora A was detected in the presence of H_2_O_2_ (Fig. 1E), or after incubation with GST alone or the Aurora A inhibitor MLN8237 (Fig. 1F).

### Identification of Cys 290 as the site of Aurora A redox regulation

The observation that oxidation-mediated inhibition of Aurora A is reversible in vitro raised the possibility that ROS such as H_2_O_2_ may serve as signaling modulators of Aurora A activity *in vivo.* The reversible oxidative modification of signaling proteins is strongly associated with the sulfur-containing amino acid cysteine (Chung et al., 2013). To provide direct evidence for Cys-SOH derivatives in Aurora A, we exploited an antibody that detects SOHs that have been selectively and covalently derivatized with dimedone (Maller et al., 2011, Seo and Carroll, 2009, Patterson et al., 2019). Sulfenate-like species were readily detected in oxidized Aurora A, and the signal increased progressively with exposure to increasing concentrations of H_2_O_2_ (Fig. 2A). Incubation of Aurora A with 10 mM DTT greatly reduced dimedone-labelling, consistent with the re-generation of Cys thiols, which are refractory to dimedone adduct formation. Importantly, no labelling of the protein was detected in the absence of dimedone, confirming antibody specificity.

**Figure 2.**
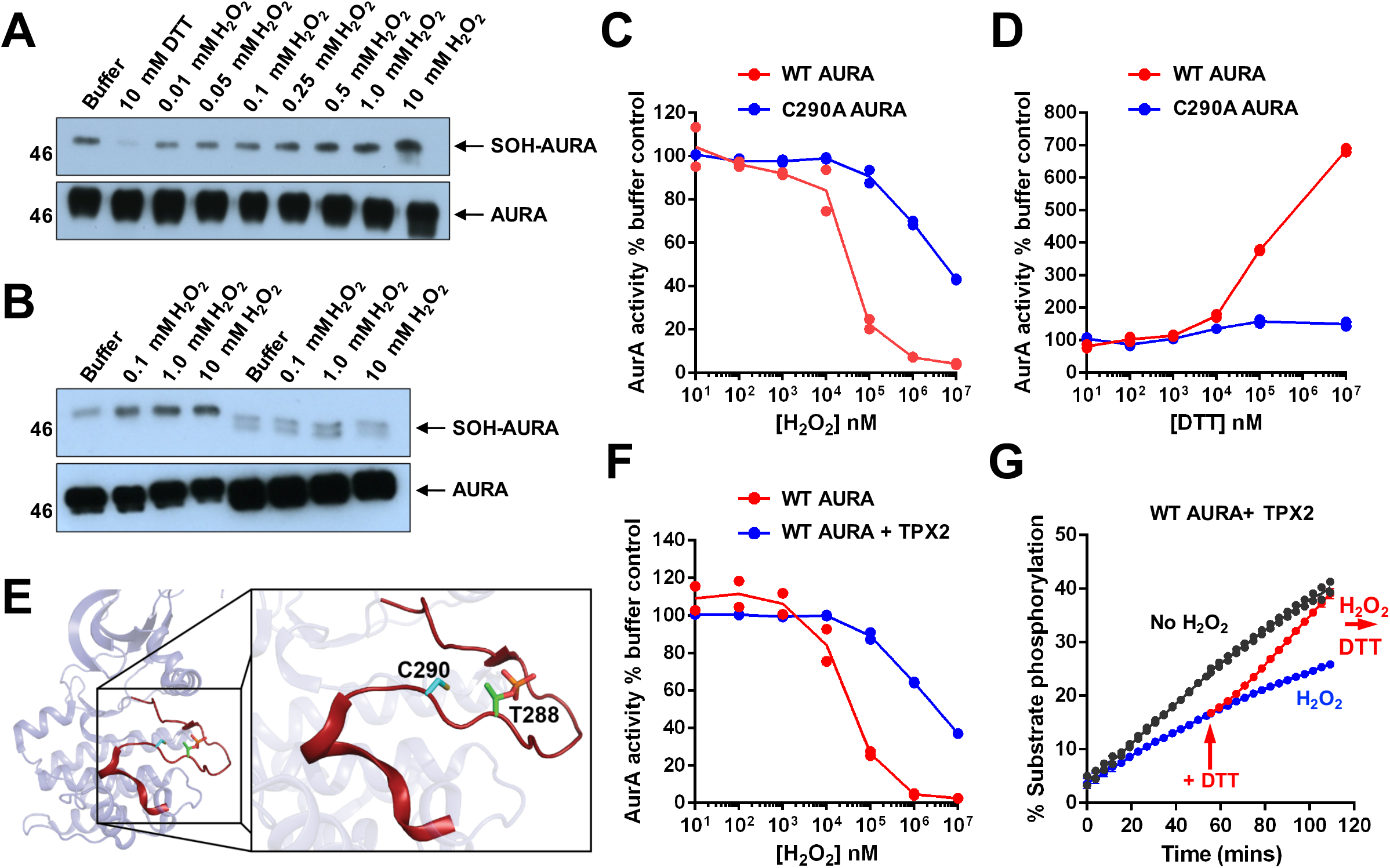
Conserved Cys 290 residue in the Aurora A activation loop is reversibly oxidized *in vitro*: effect of TPX2. **(A)** Detection of reactive cysteine oxidation in Aurora A with an antibody with specificity towards cysteine sulfenic acids that have been derivatised by dimedone (SOH-Aurora A). Aurora A (0.5 μg) was incubated with an increasing concentration of H_2_O_2_ or 10 mM DTT for 10 min and then exposed to 1 mM dimedone for a further 20 mins (all incubations performed at 20°C). Total Aurora A loading is also shown (bottom panel). **(B)** Immunoblot demonstrating depleted total SOH content in Aurora A C290A compared to WT enzyme. Assay conditions are as described for **(A). (C)** Comparative analysis of WT (red) and C290A (blue) Aurora A activity in the presence of a fixed amount of ATP (1mM) and varying concentrations of H_2_O_2_ or **(D)** DTT. Assays were performed using 6 nM Aurora A, and the extent of substrate phosphorylation (presented here as activity normalized to controls containing buffer alone) was determined after 60 mins assay time. **(E)** Structural disposition of the Aurora A activation segment (red). Cys 290 lies adjacent to Thr 288, the site of Aurora A autophosphorylation. **(F)** Binding of TPX2 protects Aurora A from inactivation by H_2_O_2_. H_2_O_2_ dose response curves are shown for Aurora A (6 nM) pre-incubated for 10 mins at 20 °C with or without 100 nM GST-TPX2 prior to oxidation with the indicated concentration of H_2_O_2_. Aurora A activity in the presence of 1 mM ATP was normalised to buffer controls after 40 mins assay time. **(G)** TPX2-Aurora A kinase activity is restored by DTT following oxidative-inhibition by H_2_O_2_. Substrate phosphorylation by Aurora A (1 nM, with 100 nM GST-TPX2) was monitored in real time in the presence or absence of 1 mM H_2_O_2_ for 50 mins prior to the addition of 2mM DTT to the indicated reactions. Assays were started simultaneously with 100 μM ATP.

To identify potential redox-sensitive Aurora A Cys residues, we employed the bioinformatics tool Cy-preds (Soylu and Marino, 2016), which consistently highlighted Cys 290 as a potential target for oxidative modification. Cys 290 lies within the canonical activation segment, in very close proximity to Thr 288 (Fig. 2E). Interestingly, the equivalent Cys in PKA has previously been analysed in terms of redox regulation, with a role for Cys 200 suggested (Humphries et al., 2005, Humphries et al., 2002). To assess whether regulatory oxidative modification of Aurora A could be assigned to Cys 290, we generated Aurora A containing a Cys to Ala substitution at this position (fig. S4A). When compared to WT Aurora A, incorporation of the C290A mutation had no effect on protein thermostability (Tm ~40 °C, fig. S4B) measured by DSF (Byrne et al., 2016, Foulkes et al., 2018) or ΔT_m_ values induced by ATP or MLN8237 binding (fig. S4C). Furthermore, K_M[ATP]_ values obtained in peptide-based kinase assays were virtually identical for both kinases, although the C290A mutant exhibited decreased activity (~50 % of WT, fig. S4D). This latter observation is consistent with previous analysis of the C290A mutation (Burgess and Bayliss, 2015), and emphasizes the potential importance of this residue as a regulatory hot-spot within the Aurora A activation loop.

We next tested whether Cys 290 was prone to oxidative modification. WT and C290A Aurora A were incubated with H_2_O_2_ to oxidize susceptible Cys residues, which were then reacted with dimedone. Immunoblotting of the derivative showed that labelling by the SOH-specific antibody was diminished in C290A Aurora A compared to WT, indicative of a reduction in the total number of Cys residues capable of undergoing reversible oxidative modification (Fig. 2B). Importantly, the mutation of Cys 290 protected Aurora A from inhibition by H_2_O_2_ (Fig. 2C). This manifested as an increase in the half-maximal inhibitory concentration (IC_50_) of H_2_O_2_ from ~35 μM for WT Aurora A to ~10 mM for C290A Aurora A. Concomitantly, C290A mutation abolished DTT dependent activation of Aurora A (Fig. 2D). Together these results confirm that Cys 290 plays a central role in a redox-regulatory mechanism that underpins Aurora A kinase activity in vitro, and suggests that this may be a direct result of a switch between oxidized ‘inactive’ and reduced ‘active’ states.

### TPX2 protects Aurora A from inactivating oxidation

In addition to activation *via* autophosphorylation of Thr 288 in the activation loop, Aurora A activity is also modulated allosterically by interactions with the spindle assembly factor TPX2 (Eyers et al., 2003, Kufer et al., 2002, Zorba et al., 2014, Dodson and Bayliss, 2012b, Ruff et al., 2018). Given that binding to TPX2 and phosphorylation of the activation loop are independent regulators of Aurora A activity, we investigated redox regulation of Aurora A in the context of TPX2 complexes. First we considered the effect of redox state on the interaction between Aurora A and TPX2. Aurora A was exposed to increasing concentrations of H_2_O_2_ or DTT and then pull-down assays were performed with GST-tagged TPX2. Aurora A remained associated with GST-TPX2 even at the highest concentrations of H_2_O_2_ and DTT employed (fig. S5A). Thus, the redox state of Aurora A had no detectable effect on binding to TPX2 in vitro. Next, we examined inhibition of Aurora A by H_2_O_2_ in the presence of TPX2. The phosphorylated activation loop of Aurora A has recently been shown to adopt a range of conformations in solution, only becoming highly ordered in a stable DFG-In conformation upon TPX2 binding (Ruff et al., 2018). Furthermore, both ‘inactive’ unphosphorylated and ‘active’ phosphorylated Aurora A adopt similarly well-defined structures upon TPX2 binding, resulting in an increase in kinase activity (Bayliss et al., 2003, Ruff et al., 2018). We found that inhibition of Aurora A by H_2_O_2_ was prevented by TPX2, increasing the IC_50_ [H_2_O_2_] value to > 1 mM (Fig. 2F). Importantly, the modest inhibitory effect of H_2_O_2_ for TPX2-Aurora A activity could be completely reversed upon addition of DTT (Fig. 2G). Allosteric activation of Aurora A by TPX2 is therefore sufficient to prevent kinase inactivation by oxidation. It is possible that oxidation of Aurora A at Cys 290 alters the structural dynamics of the activation loop and stabilizes a less active subpopulation, with activity being recapitulated following binding to- and structural reorganization by TPX2. This is supported by the observation that C290A Aurora A, which displays weaker kinase activity compared to the WT protein, is also strongly activated by TPX2 binding, but unlike WT Aurora A was completely resistant to inhibition by H_2_O_2_ when in a TPX2-bound state (fig. S5B, C). Importantly the activities of both WT and C290A Aurora A were unaffected by GST (fig. S5B).

### Aurora A can be activated by glutathionylation on Cys290

The glutathionylation of kinases such as PKA can lead to changes in protein function (Humphries et al., 2002). We therefore investigated the influence of reduced (GSH) or oxidized (GSSG) glutathione alongside a panel of other redox-active compounds on Aurora A activity. Aurora A activation by the reducing agents DTT, tris(2-carboxyethyl)phosphine (TCEP) and 2-mercaptoethanol (2-ME) was confirmed by enhanced rates of substrate phosphorylation compared to control assays (Fig. 3A). Rather surprisingly, the inclusion of GSH (or GSSG) in the assay also induced a measurable increase in activity (Fig. 3A). This is in contrast to the redox-regulated kinase PKA, which is reported to be inhibited by GSH (Humphries et al., 2002). Consistently, C290A Aurora A was resistant to activation by reducing agents (Fig. 3B). Importantly, and in contrast to WT Aurora A, which demonstrated concentration-dependent activation by GSH, C290A Aurora A only exhibited modest changes in activity at the highest tested concentration of GSH (Fig. 3C). Next we investigated whether modulation of Aurora A activity was a consequence of mixed disulfides forming between Cys 290 and glutathione. To probe for glutathionylation, we employed an antibody that specifically reacts to protein-glutathione complexes (Fig. 3D). Glutathionylation was readily detected for WT Aurora A incubated with GSSG, but not for the C290A mutant, suggesting that changes in Aurora A activity were a direct result of disulfide exchange with the metabolite (Fig. 3D). Furthermore, glutathionylated Aurora A was completely eliminated under reducing SDS-PAGE conditions, confirming the presence of a disulfide bond (fig. S6A). PKA was included as a positive control as it has previously been shown to be modified and inactivated by glutathione (Humphries et al., 2002). However, despite stimulating Aurora A activity, addition of GSH alone was insufficient to restore the activity of oxidized Aurora A (Fig. 3E), which required reduction by DTT (fig. S6B) or enzymatic deglutathionylation by glutaredoxin-1 (GRX) (fig. S6C) to restore Aurora A catalytic activity. The AGC kinase AKT has previously shown to be regulated by glutathione-dependent mechanisms (Murata et al., 2003b). To evaluate AKT glutathionylation using a real-time assay, PDK1-phosphorylated S473D AKT was incubated with GSH in the presence and absence of H_2_O_2_. Similar to Aurora A, AKT was covalently modified by glutathione (Fig. 3F). The catalytic activity of AKT was also enhanced hundreds-of-fold by exposure to GSH in the absence of H_2_O_2_ (fig. S6D). Furthermore, the activity of oxidized AKT was inhibited by oxidation, and could be restored by the addition of DTT; however, the addition of GSH had no effect (Fig. 3G), as demonstrated for Aurora A.

**Figure 3.**
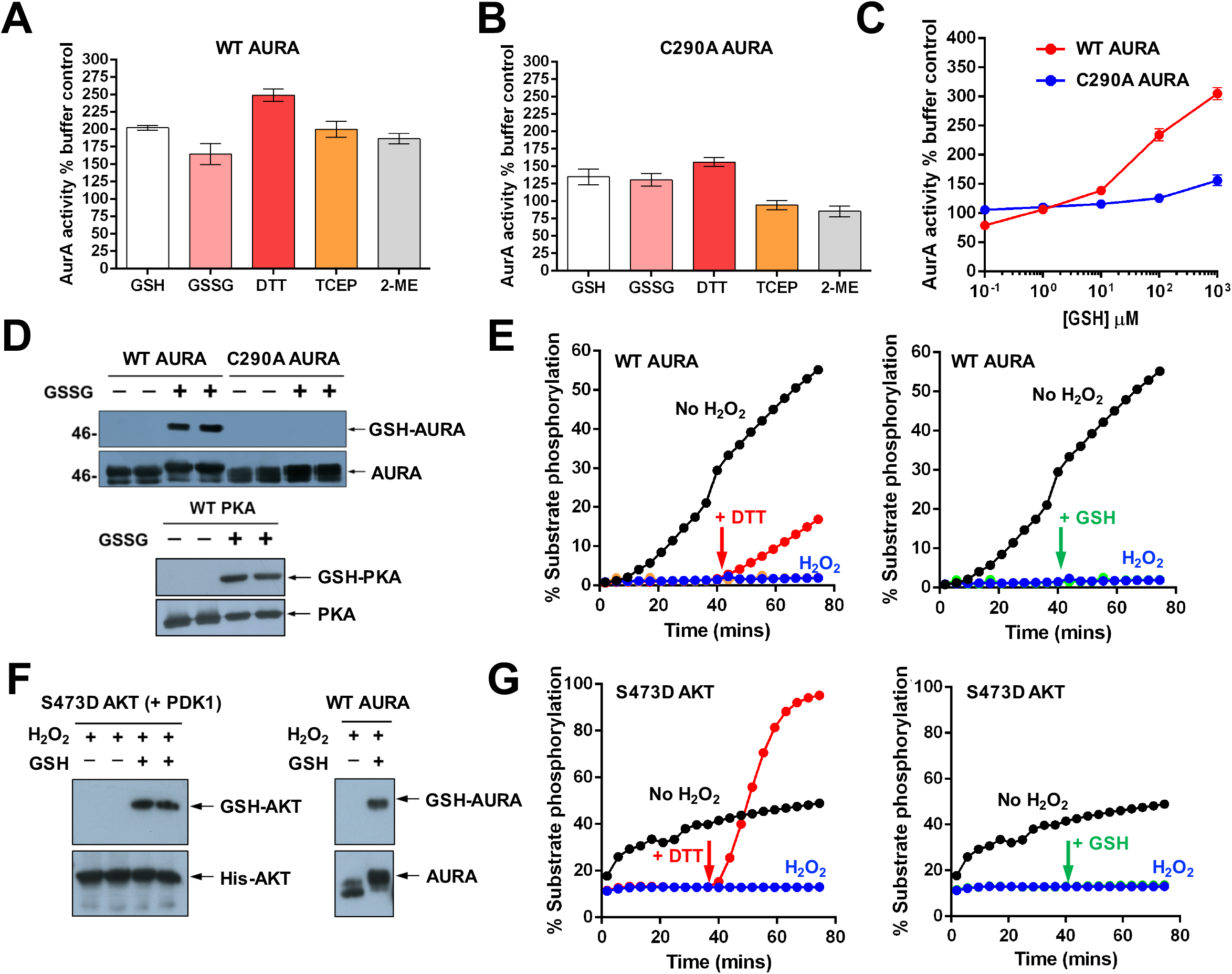
Aurora A is activated by modification of Cys 290. **(A)** Redox-dependent activation of Aurora A or **(B)** C290A Aurora A by a panel of reducing agents. WT and C290A Aurora A proteins (12.5 nM) were incubated with 1 mM ATP in the presence of 1 mM of the appropriate reducing agent and fluorescent peptide substrate. Activity was calculated relative to a control (buffer only) after 40 mins assay time. **(C)** Dose response curves for GSH. The activity of WT and C290A Aurora A (12.5 nM) was monitered in the presence of increasing concentrations of GSH and 1 mM ATP. Aurora A activity was normalised to buffer controls after 40 min assay time. **(D)** Immunoblot demonstrating in vitro glutathionylation of Aurora A at Cys 290. 1 μg of recombinant purified WT and C290A Aurora A proteins were incubated in the presence or absence of 10 mM GSSG for 30 mins at 20°C. Western blots were probed with an antibody with specificity towards glutathione-conjugated proteins. Equal loading of protein was confirmed using an antibody for Aurora A. PKA (1 μg) was also included as a positive control. SDS-PAGE was performed under nonreducing conditions. **(E)** Inhibition of Aurora A by H_2_O_2_ is relieved by DTT but not GSH. Aurora A (12.5 nM) activity was assayed in real time in the presence or absence of 1 mM H_2_O_2_ for 50 mins and reactions were supplemented (where indicated) with 2 mM DTT (left panel) or GSH (right panel). Aurora A-dependent phosphorylation of the fluorescent peptide substrate was initiated by the addition of 1 mM ATP. **(F)** Immunoblot demonstrating glutathionylation of AKT. 1 μg PDK1 phosphorylated S473D AKT was incubated with 100 μM H_2_O_2_ for 10 mins prior to the application of 1 mM GSH (30 mins at 20°C). SDS-PAGE was performed under non-reducing conditions and 1 μg Aurora A was used as a positive control. **(G)** Activity of AKT is recapitulated by DTT but not GSH. The activity of PDK1 phosphorylated S473D AKT (7 nM) with 1 mM ATP was monitored in the presence of 1 mM H_2_O_2_ for 40 mins prior to the addition of 2mM DTT (left panel) or GSH (right panel).

### Oxidative stress inhibits Aurora A substrate phosphorylation in human cells

Next, we investigated redox regulation of Aurora A in cells, employing endogenous TACC3 phosphorylation as an intracellular marker for active Aurora A (Kinoshita et al., 2005, Tyler et al., 2007). HeLa cells were initially synchronized with nocodazole, and then exposed to H_2_O_2_ or DTT for 30 mins. Western blotting revealed a marked decrease in TACC3 phosphorylation at Ser 558 in cells treated with H_2_O_2_ compared to control cells or those treated with DTT (Fig. 4A). Consistently, cells incubated with the Aurora A inhibitor MLN8237, demonstrated complete loss of TACC3 phosphorylation (Fig. 4A), and nocodazole alone, which has recently been shown to enhance the oxidation of Cys residues (Patterson et al., 2019), did not lead to general Aurora A inhibition. Aurora A protein levels were not affected by the presence of H_2_O_2_, suggesting that loss of TACC3 phosphorylation was due to inhibition of Aurora A rather than destabilization of the protein. The extent of TACC3 phosphorylation was also decreased by H_2_O_2_ in a dose-dependent manner (Fig. 4B). We next determined the effects of the cell-permeable oxidant diamide (Kosower and Kosower, 1995). Strikingly, the extent of TACC3 phosphorylation was inversely proportional to the diamide concentration, analogous to observations with H_2_O_2_ (Fig. 4C). Menadione is a quinone oxidant that stimulates rapid generation of cellular ROS through redox-cycling (Criddle et al., 2006). Menadione exposure also caused a concentration-dependent inhibition of TACC3 phosphorylation (Fig. 4D). Finally, we investigated the effect of chronic oxidative stress on Aurora A activity. Glucose oxidase (GO) was added to nocodazole-containing culture medium at a non-toxic concentration (2 U/ml) to facilitate the generation of peroxide, inducing intracellular steady-state levels of H_2_O_2_ of 1-2 μM (Askoxylakis et al., 2011, Mueller et al., 2009, Truong et al., 2016). As shown in Fig. 4E, GO exposure also resulted in a time-dependent decrease in TACC3 phosphorylation.

**Figure 4.**
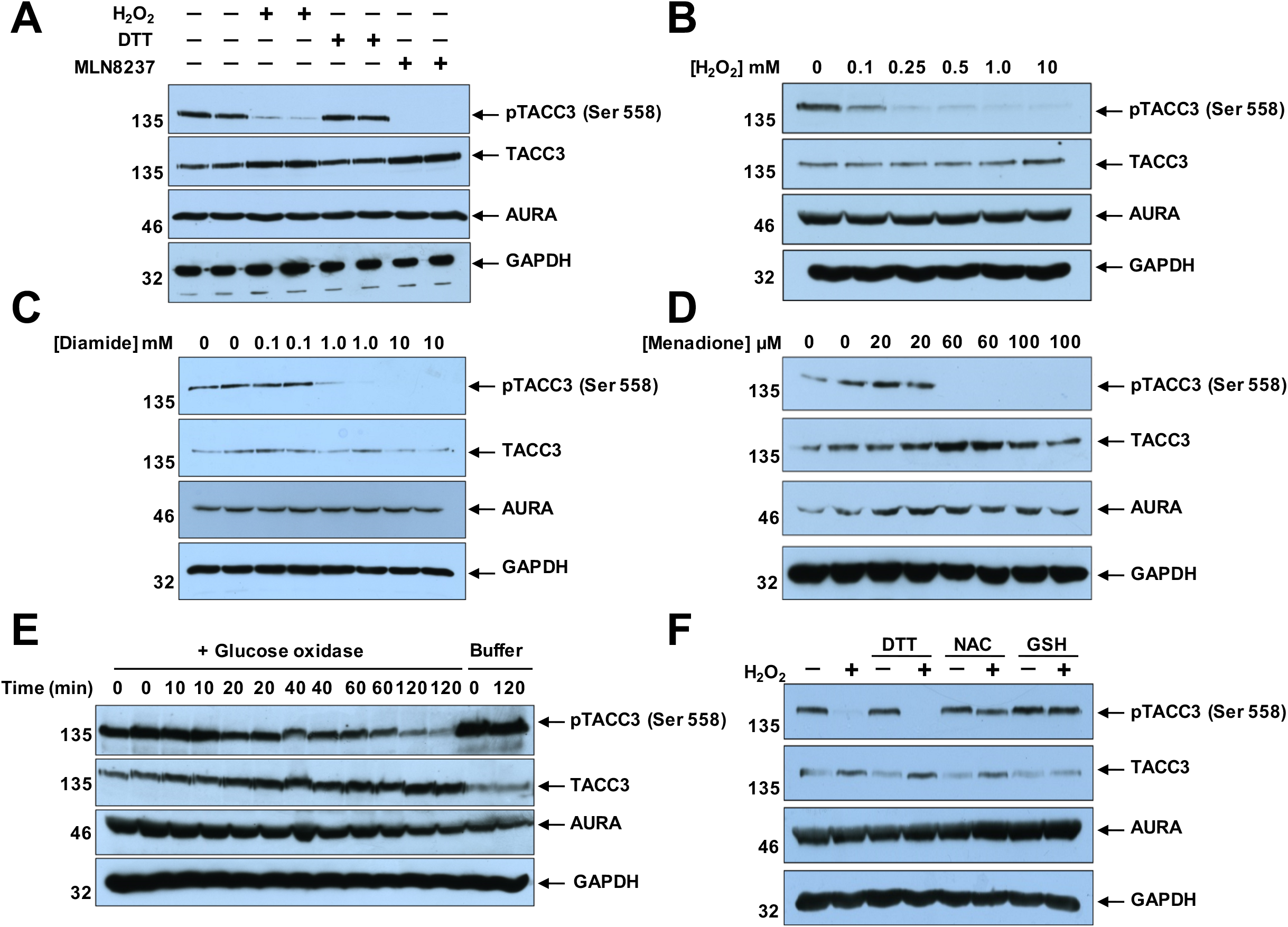
Oxidation inhibits endogenous Aurora A activity in human cells. **(A)** Treatment of HeLa cells with H_2_O_2_ results in a loss of TACC3 phosphorylation at Ser558. Immunoblot shows a loss of Aurora A-dependent phosphorylation of TACC3 (pTACC3) in HeLa cells untreated or treated with 10 mM H_2_O_2_ or 1 μM MLN8237 for 30 mins. **(B)** Concentration dependent inhibition of endogenous Aurora A activity by H_2_O_2_. Immunoblots show a loss of pTACC3 in HeLa cells incubated with the indicated concentration of H_2_O_2_ for 30 mins. A representative western blot of two independent experiments is shown. **(C)** Concentration dependent inhibition of endogenous Aurora A activity by the oxidizing reagent diamide and redox-cycling quinone menadione. **(D)** Loss of pTACC3 in HeLa cells incubated with the indicated concentration of diamide for 30 mins or menadione for 60 mins. **(E)** Loss of pTACC3 in HeLa cells exposed to glucose oxidase (GO). Immunoblots show time-dependent inhibition of TACC3 phosphorylation in HeLa cells cultured in the presence of 2 U/ml GO and nocodazole for the indicated time periods. **(F)** Aurora A oxidative-inhibition is reversed by the peroxide scavengers, NAC and GSH. Immunoblots of HeLa cells treated with 10 mM H_2_O_2_ for 10 mins prior to the addition of fresh culture medium containing 10 mM DTT, NAC or GSH or buffer control. Cells were then cultured for an additional 20 mins prior to the extraction of whole cell lysates. In all experiments described here, HeLa cells had been blocked in mitosis by nocodazole (100 ng/mL) for 16 h. Whole cell lysates were analysed by western blot and probed with anti-TACC3 or anti-pSer558-TACC3 antibodies. Equal loading was confirmed using antibodies against Aurora A and GAPDH.

To evaluate the potential physiological relevance of Aurora A redox regulation as a signaling mechanism, we investigated the reversibility of inactivation. Cells were exposed to H_2_O_2_ prior to incubation with DTT or the cellular antioxidants GSH and N-acetyl-L-cysteine (NAC). Under these conditions, TACC3 phosphorylation was restored to basal levels by both GSH and NAC, presumably due to ROS scavenging (Fig. 4F), and consistent with our in vitro rescue data (fig. S6C). In the presence of H_2_O_2_, DTT did not rescue TACC3 phosphorylation. Taken together, these results demonstrate, for the first time, reversible redox-dependent regulation of Aurora A in cells, confirming our biochemical analysis.

### Cys residues are evolutionary conserved in ePK activation segments

Cys is the second least-common amino acid in vertebrate proteomes (van der Reest et al., 2018), although >200,000 Cys residues are present in the human proteome (Go et al., 2015), and reactive Cys side chains, especially those lying on surface-exposed regions of proteins (Soylu and Marino, 2016), are susceptible to redox modification (fig. S1). To investigate the potential generality of Cys-based redox mechanism in protein kinases that possess an activation segment, we analyzed >250,000 protein kinase related sequences, confirming that ∼11.5% of ePKs found across the kingdoms of life possess the Cys 290 equivalent of Aurora A (Fig. 5A). This number further reduced to 1.4% of all ePKs when the co-conservation of Cys residues at the DFG +2 and ‘T-loop +2’ positions, were considered together (Fig. 5A, bottom). Remarkably, the Cys 290-equivalent residue in the 97 human kinases (Table 1) was very strongly associated with two of the seven human kinase groups, the well-studied AGC kinases and CAMKs (Fig. 5B), and this pattern was also observed across the kingdoms of life (fig. S7A). To evaluate this observation experimentally, we purified and analysed a variety of Cys-containing protein kinases (Fig. 5C), beginning with PKA, PLK1, PLK4 and MELK, which were purified from bacteria in the absence of DTT (fig. S8) and assayed using a redox microfluidic kinase assay established for Aurora A and using kinase-specific peptide substrates (Table 2). The activity of each kinase was measured in the presence or absence of increasing concentrations of H_2_O_2_ or DTT and the reversibility of redox-dependent modulation confirmed using a H_2_O_2_ and DTT rescue procedure. In agreement with published findings, PKA activity was inhibited in a concentration dependent manner by H_2_O_2_ (Humphries et al., 2005, Humphries et al., 2007a, Humphries et al., 2002) whereas DTT modestly stimulated kinase activity (Fig. 6A). Importantly, the inhibitory effect of H_2_O_2_ was completely abrogated in the site-specific cysteine mutant PKA C200A (Fig. 6A), in agreement with previous findings (Humphries et al., 2002). Inhibition of PKA by H_2_O_2_ was due to the reversible oxidation of a sulfhydryl residue, since activity could be partially restored by DTT (Fig. 6B).

**Figure 5.**
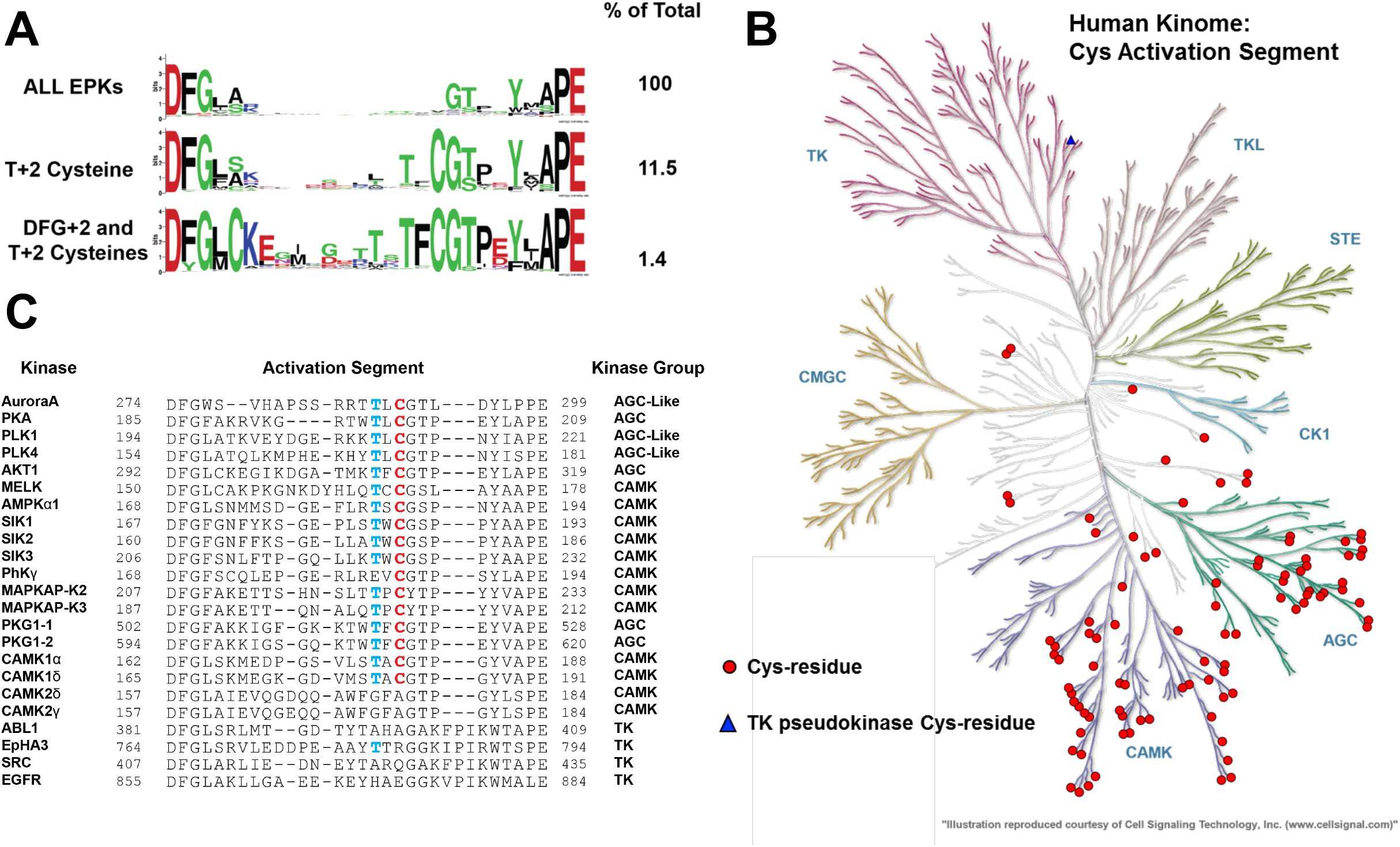
Bioinformatic analysis of Aurora A Cys 290-equivalent in all ePKs. **(A)** Analysis of ePKs, centered on the activation segment between the canonical DFG and APE motifs. The amino acid distribution (percentage of all kinases) is shown on right, data presented as HMM Sequence Logos. **(B)** Human Kinome dendrogram, showing highly skewed distribution of kinases containing a Cys residue at the T-loop +2 residue in AGC and CAML groups. **(C)** Activation segment alignment of human kinases analysed in this study.

**Figure 6.**
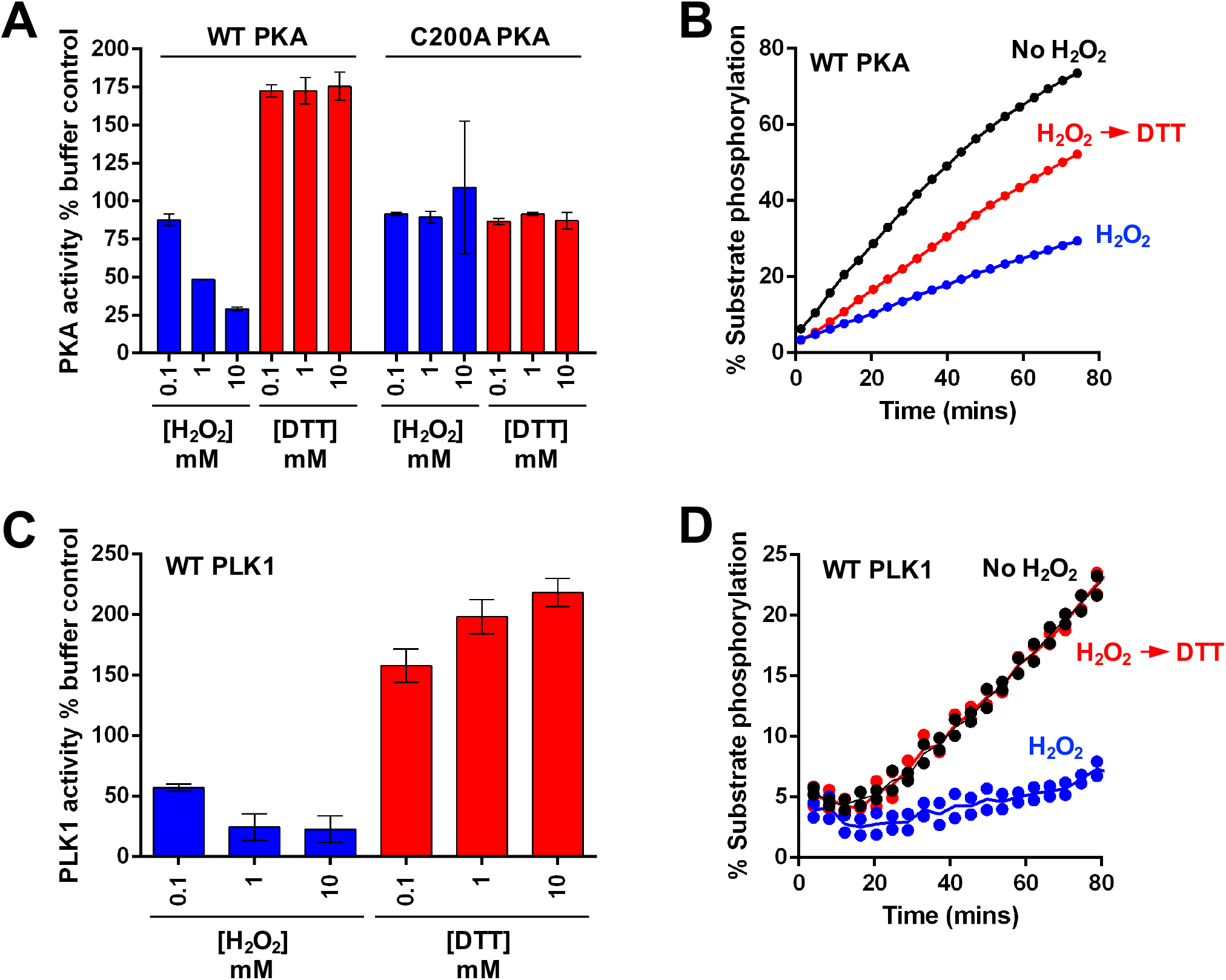
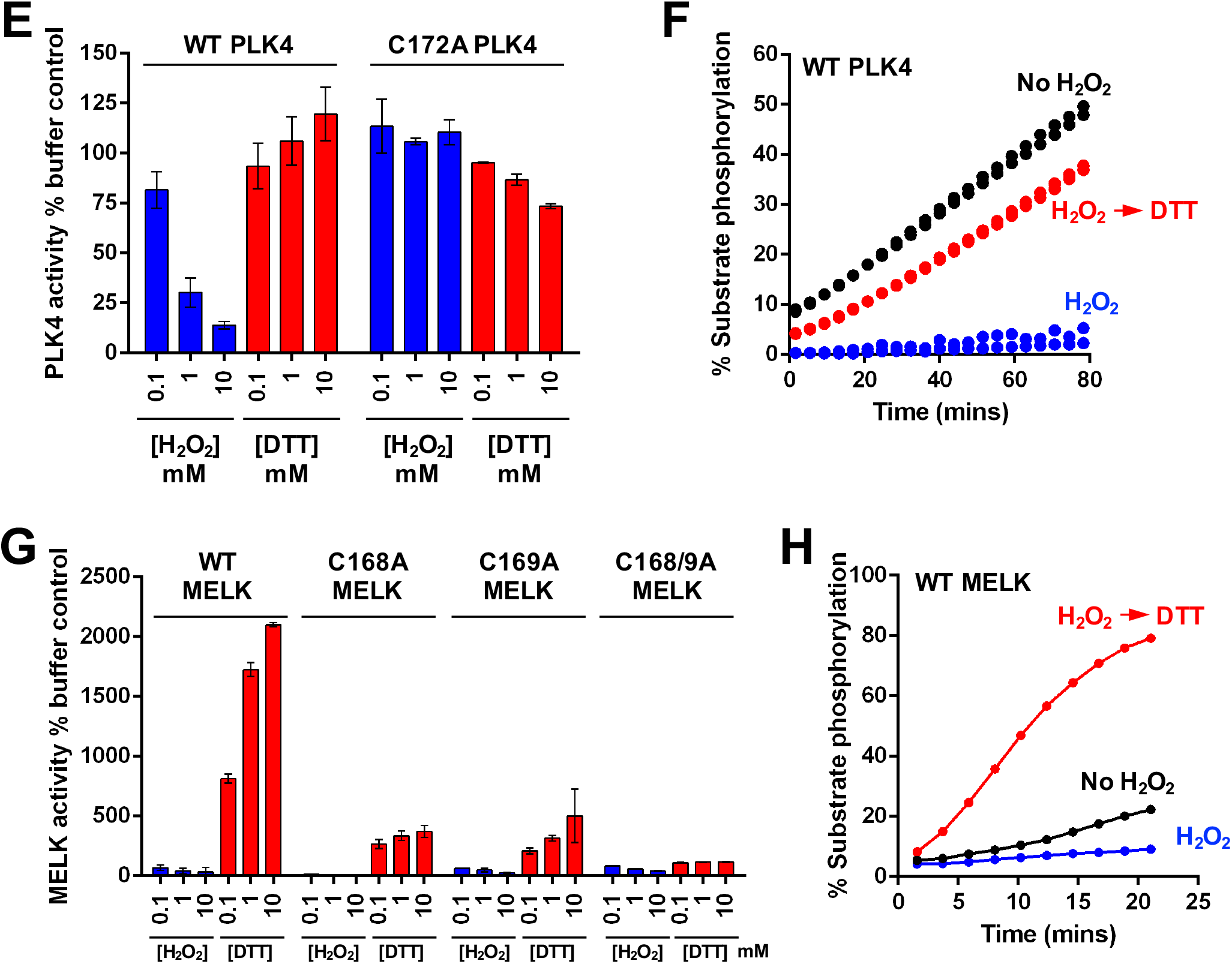
Reversible oxidation of a conserved Cys residue regulates the activities of many Ser/Thr kinases. **(A)** Redox regulation of PKA catalytic domain. WT or C200A His-PKA (0.3 nM) were assayed with the indicated concentration of H_2_O_2_ or DTT and activities were normalized relative to controls. **(B)** Reversible redox regulation of PKA. **(C)** Redox regulation of PLK1 catalytic domain. GST-PLK1 (160 nM) was assayed as for PKA, in the presence of the indicated concentration of H_2_O_2_ or DTT and activity was normalized relative to buffer control. **(D)** Reversible redox regulation of PLK1. **(E)** Redox regulation of WT and C172A His-PLK4. His-PLK4 (4 μM) was assayed with H_2_O_2_ or DTT and activity was normalized relative to control. **(F)** Reversible redox regulation of WT PLK4. **(G)** Redox regulation of MELK. WT and activation loop Cys mutants (50 nM) were assayed with H_2_O_2_ or DTT and activities normalized relative to controls. **(H)** Reversible redox regulation of WT MELK. In these assays, kinases were incubated on ice for 30 mins in the presence or absence of 5 mM H_2_O_2_. Reactions were then initiated with the addition 1 mM ATP and substrate peptide in the presence or absence of 10 mM DTT. Protein concentrations were as previously described. For PKA assays, H_2_O_2_ and DTT were used at 10 and 20 mM respectively. In all experiments, kinase-optimised fluorescent peptide substrates were employed (see Table 2).

**Table 1.**
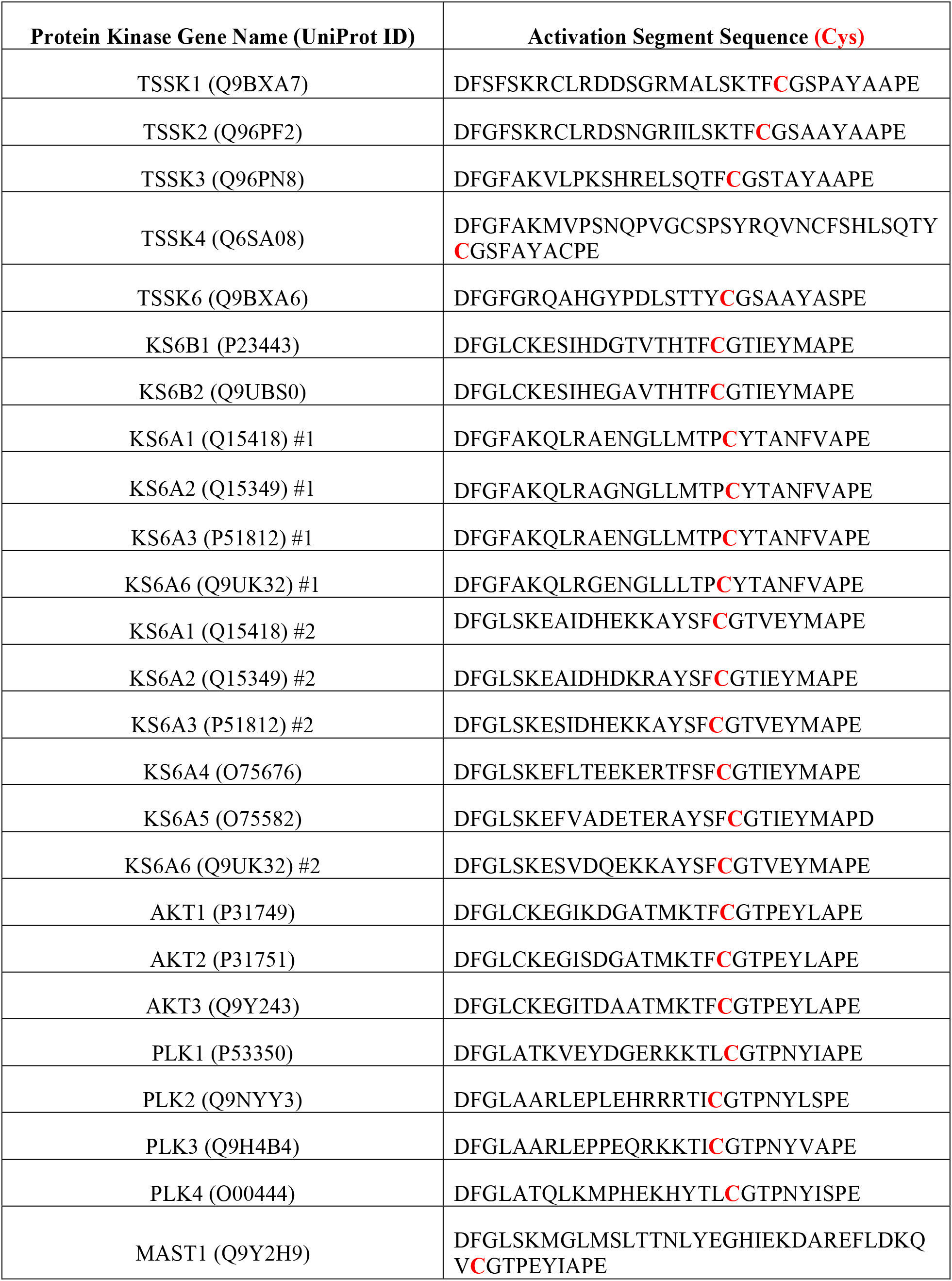
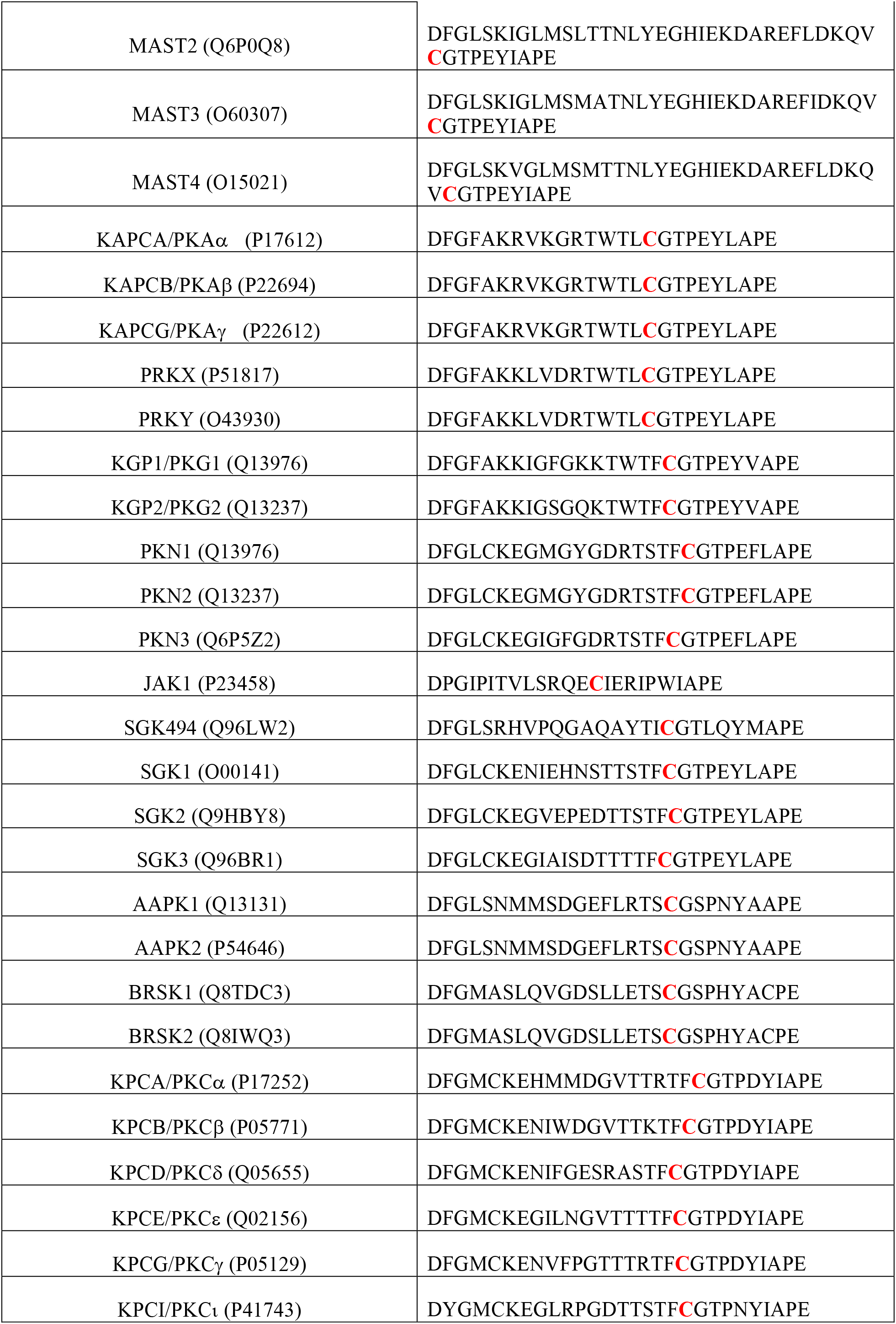
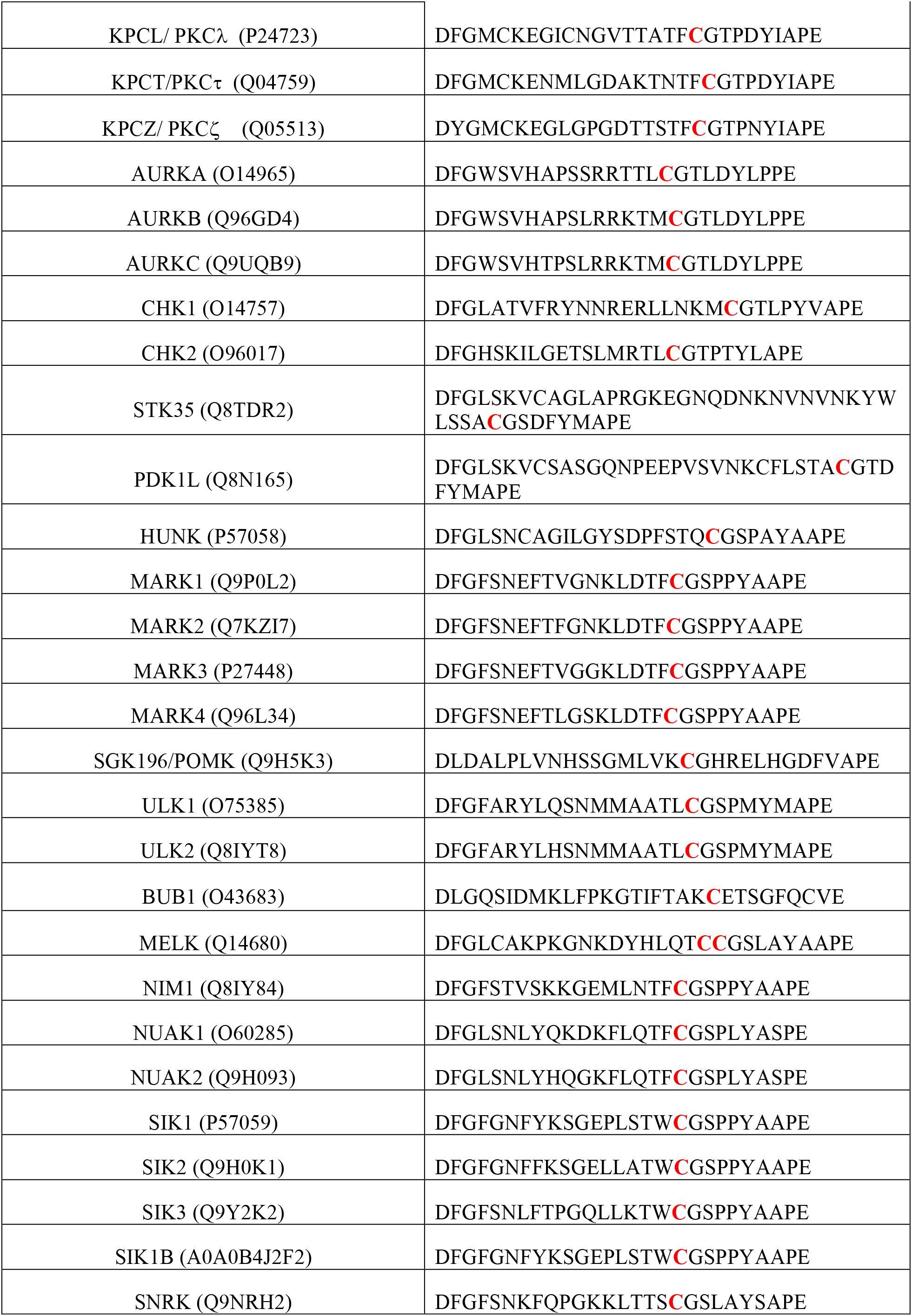
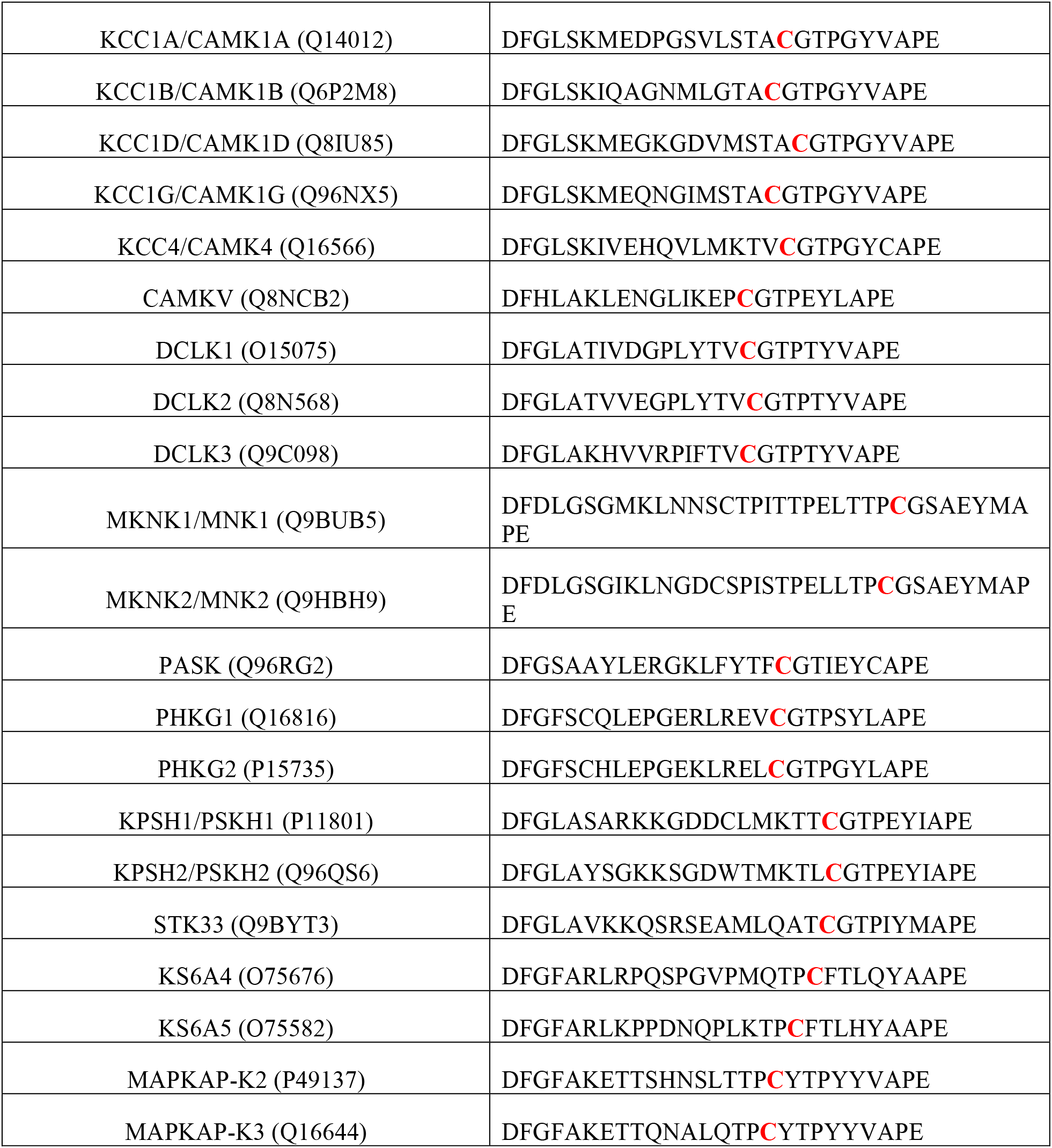
Human protein kinases containing the Aurora A Cys 290 equivalent in the Activation Segment

**Table 2.**
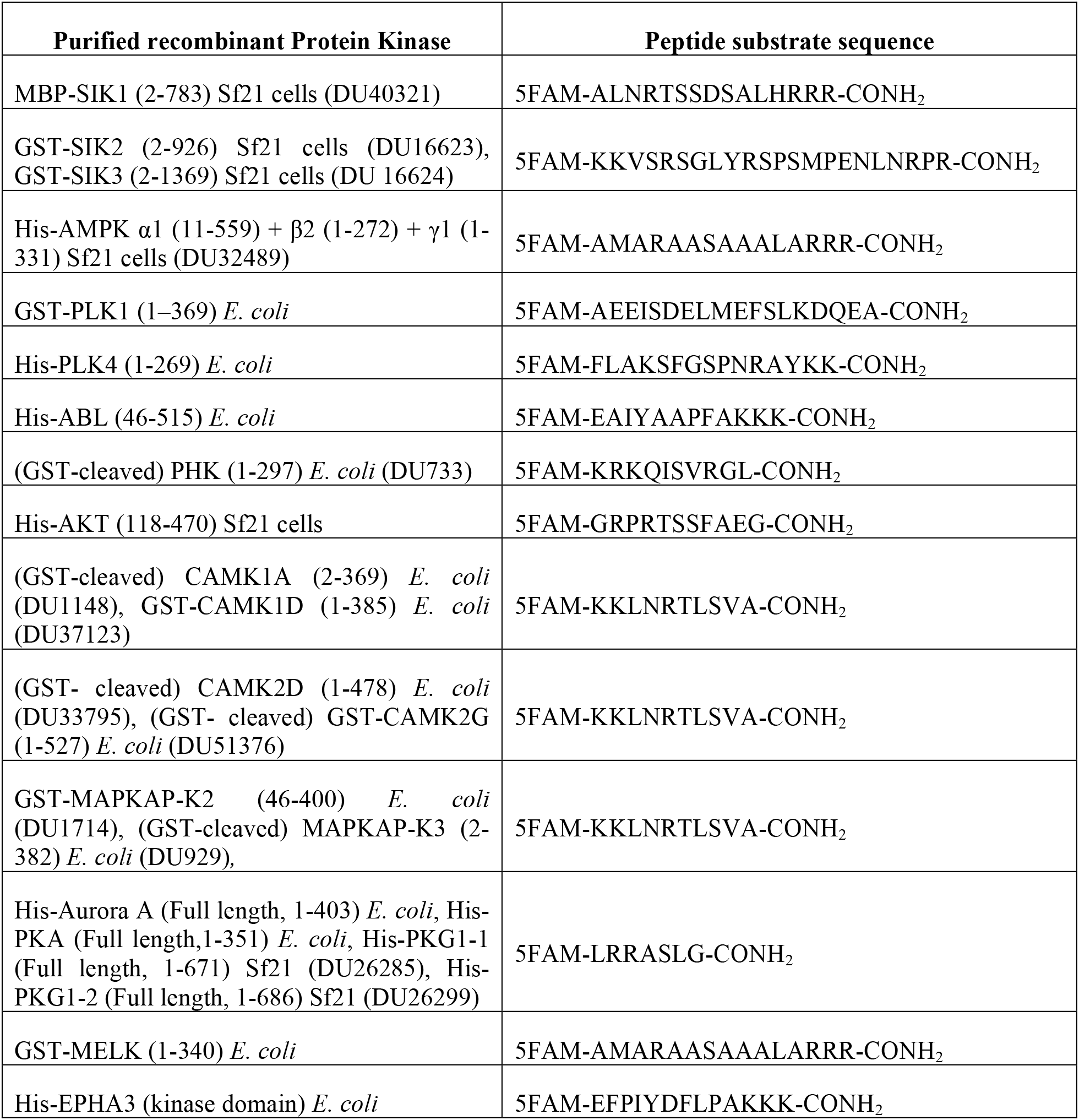
Protein kinase enzymes and substrates. Sequence of recombinant protein kinases and peptide substrates employed for assay of purified human protein kinases. The sources of the enzymes (insect cell or bacteria) are also included. Statistical analysis. All experimental procedures were repeated in at least 3 separate experiments with matched positive and negative controls (unless stated otherwise). Results are expressed as mean ± SD for all in vitro experiments and data are expressed as the mean ± standard deviation. When applied, statistical significance of differences (*P ≤0.05) was assessed using a Students t-test for normally-distributed data. All statistical tests were performed using Prism 7 (GraphPad Software).

### Cys-dependent redox regulation of Aurora A-related kinases

The Aurora kinases and Polo-like kinases (PLKs) both perform related mitotic functions (Leroux et al., 2018). The PLK family are stringently regulated multifaceted modulators of mitosis and cytokinesis (Combes et al., 2017), and the close regulatory relationship between PLK1 and Aurora A in mitosis (Macurek et al., 2008, Scutt et al., 2009) led us to investigate potential redox regulation for both PLK1 and PLK4. (Caron et al., 2016a). We demonstrated dose-dependent activation and inhibition of PLK1 by DTT and H_2_O_2_ respectively (Fig. 6C), and inhibitory oxidation was completely reversed by subsequent exposure to DTT (Fig. 6D). Mutation of the conserved Cys in the PLK1 activation segment (Cys 172) abolished activity, suggesting an important role for this residue in attaining an active fold. Next, we evaluated PLK4, the master centrosomal kinase (Bettencourt-Dias et al., 2005, Habedanck et al., 2005), which also possess a Cys residue (Cys 172), +2 residues from the T-loop autophosphorylation site (Thr 170). PLK4 was reversibly inhibited by oxidation and stimulated under reducing conditions (Fig. 6E & F). C172A PLK4 was totally refractory to inhibition by oxidation, confirming the role of this conserved cysteine as a redox-dependent regulator of activity in this kinase (Fig. 6E). An evolutionary analysis of all kinases also clearly demonstrated that the Cys 290-equivalent was conserved across eukaryotic Aurora, PKA and Polo-like kinases (fig. S7B-E)

Maternal embryonic leucine zipper kinase (MELK) is a member of the CAMK kinase grouping, and related to the AMPK-related kinases (Manning et al., 2002, Wang et al., 2014). Redox regulation of MELK has previously been reported, although the precise mechanism remains unclear (Beullens et al., 2005). Interestingly, the activation segment of MELK contains two consecutive Cys residues (Fig. 5C), one of which is predicted to form an intermolecular disulfide bond with an equivalent Cys supplied by a dimeric partner (see Discussion). MELK exhibited very low activity in the absence of DTT, with activity increasing several hundred-fold after inclusion of DTT in the assay (Fig. 6G). Interestingly, H_2_O_2_ inhibited MELK activity in a dose-dependent manner (Fig. 6G), and DTT-dependent activation was so pronounced that MELK activity rapidly surpassed control levels when DTT was used to ‘rescue’ H_2_O_2_-inhibition (Fig. 6H). Both C168A and C169A point mutation blocked DTT-dependent MELK activation, but neither mutation in isolation completely abrogated MELK redox-sensitivity (Fig. 6G). However, combined mutation (C168A/C169A MELK) completely abolished redox regulation, and Cys mutations diminished MELK oxidation, and dimedone adduct formation, particularly under non-oxidizing conditions (fig. S8B), extending previous findings (Beullens et al., 2005).

### Evaluation of redox regulation in a Cys-containing CAMK and AGC kinase panel

To establish the generality of our findings, we increased the scope of our analysis to incorporate a panel of protein kinases that contain an evolutionary-conserved Cys residue in the T-loop +2 position of the activation segment (Fig. 5C). Purified enzymes were supplied in the presence of 1 mM DTT, and assayed using specific peptide substrates. Catalytic activity was quantified in the presence of additional DTT, H_2_O_2_ or H_2_O_2_ ‘rescued’ with DTT. Remarkably, the majority of kinases tested displayed redox-dependent regulation. We previously showed that AKT was activated by GSH (fig. S6D) and AKT outputs in cells are redox-sensitive (Durgadoss et al., 2012, Tan et al., 2015). We confirmed that DTT enhances AKT activity several hundred-fold (Fig. 7A). In contrast, oxidation completely prevented AKT-dependent substrate phosphorylation, although activity was restored, or even enhanced, by subsequent DTT exposure (Fig. 7A). The 5′-AMP-activated protein kinase holoenzyme complex (AMPK, comprised of α1, β2 and γ1 subunits) is a member of the CAMK family and was also strongly inhibited by H_2_O_2_ in a DTT-reversible manner (Fig. 7B). This finding supports a growing body of evidence that suggests ROS participate in the regulation of AKT and AMPK activity, although the precise mechanisms remain controversial. Direct activation of AMPKα catalytic subunits in the presence of H_2_O_2_ has previously been reported (Zmijewski et al., 2010), whereas inhibition of AMPKα activity through Cys130/174 oxidation has also previously been shown (Shao et al., 2014). Cys174 in AMPKα is analogous to Cys 290 of Aurora A, and is situated upstream of the Thr172 phosphorylation site, a critical modulator of AMPK activity.

**Figure 7.**
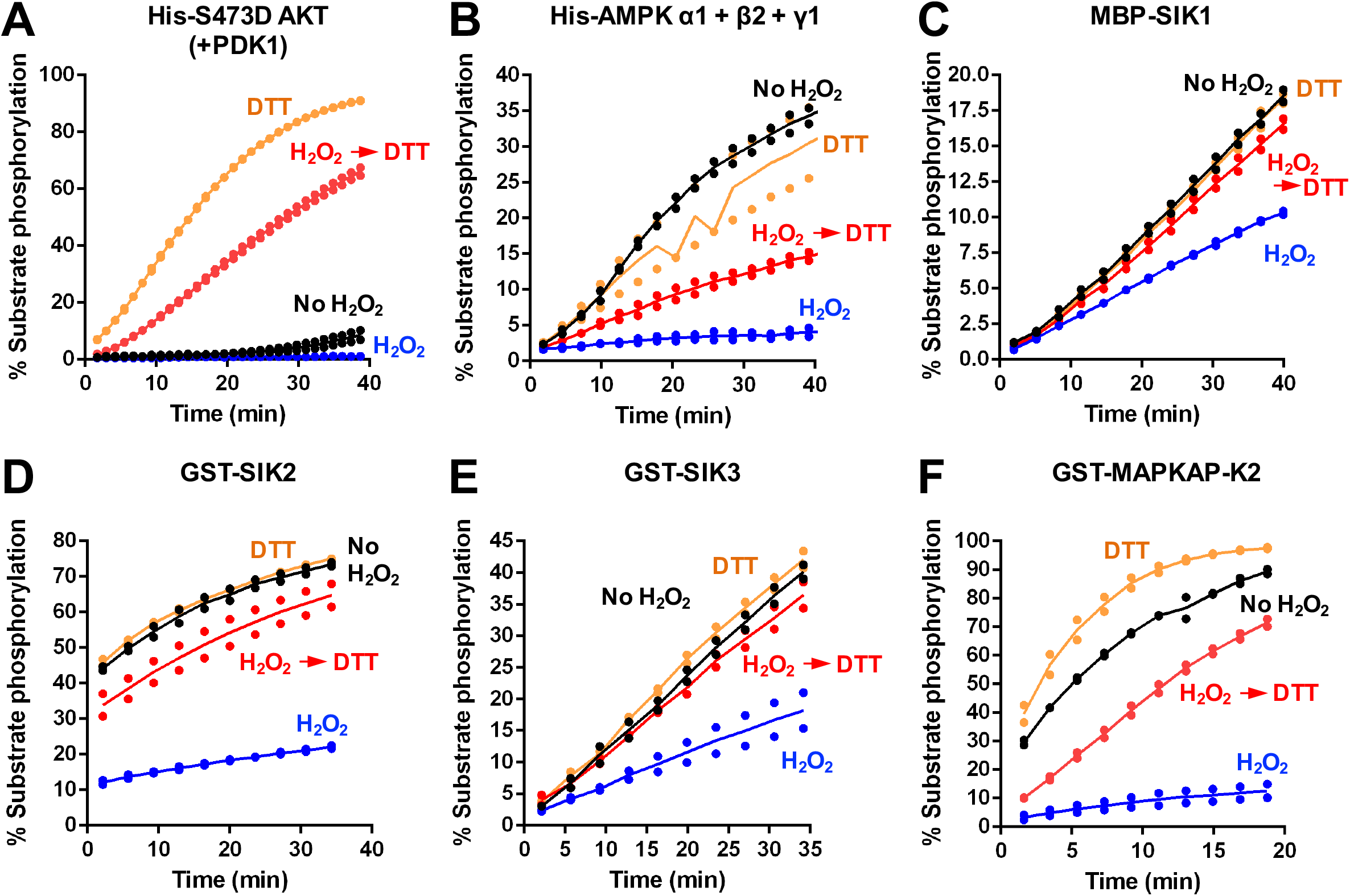
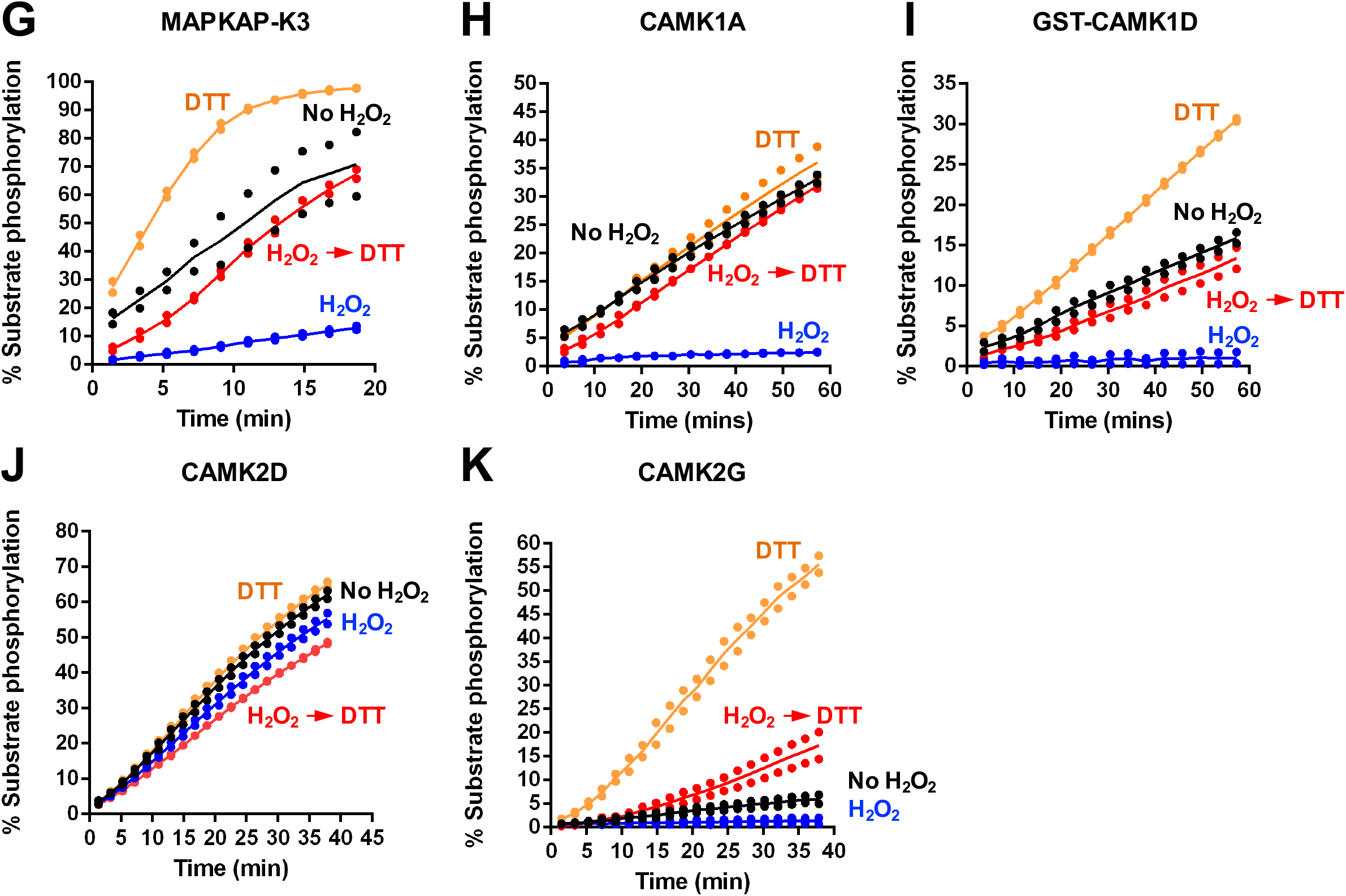
T-loop +2 Cys residue is partially diagnostic for reversible redox modulation of catalytic output in human AGC and CAM kinases. A cohort of AGC and CAMK-related kinases were probed for reversible oxidation-dependent inhibition using real time phosphorylation of kinase specific peptide substrates (Table 2). In all assays, kinases were incubated on ice for 30 mins in the presence or absence of 5 mM H_2_O_2_. Reactions were then initiated with the addition 1 mM ATP and substrate peptide in the presence (where indicated) of 10 mM DTT. Kinases were assayed at the following concentrations; **(A)** 7 nM PDK1 phosphorylated His-S473D AKT, **(B)** 24 ng of His-AMPK a1 + ß2 + γ1, **(C)** 0.5 μM MBP-SIK1, **(D)** 2 nM GST-SIK2, **(E)** 2 nM GST-SIK3, **(F)** 10 nM GST-MAPKAP-K2, **(G)** 15 nM MAPKAP-K3, **(H)** 0.7 μM CaMK1A, **(I)** 60 nM GST-CaMK1D, **(J)** 0.6 μM CaMK2D and (K) 0.6 μMCaMK2G. Assays including CaMK were supplemented with 5 mM CaCl_2_ and 2 μM Calmodulin.

The salt-inducible kinases (SIK1-3) are also members of the AMPK-related family of CAMKs (Bright et al., 2009). Purified SIK1-3 were also reversibly inhibited by H_2_O_2_ (Fig. 7C-E). Interestingly, H_2_O_2_ was also weakly- and reversibly inhibitory towards phosphorylase kinase, PHKγ, a founding member of the CAMK group (fig. S9A). We next analyzed redox regulation in the CAMK-family members MAPKAP-K2 and MAPKAP-K3, demonstrating potent inhibition of both enzymes by peroxide (Fig. 7F-G). In contrast, DTT activated both kinases, and also restored activity following peroxide treatment (Fig. 7F-G). We ruled out an effect of kinase redox modulation through an indirect effect of GST, since the tag was absent in the MAPKAP-K3 preparation, but present in GST-MAPKAP-K2.

Protein Kinase G (PKG) belongs to the AGC group of kinases, although its mechanism of regulation is distinct from that of closely related PKA and PKC isozymes. Moreover, in contrast to other T-loop Cys-containing AGC kinases tested (including PKA and AKT) there was no evidence for oxidative inhibition of either of the PKG1 splice variants, PKG1-1 or PKG1-2, when they were assayed in either the absence, or presence, of cGMP (fig. S9B-E), consistent with recent findings (Kalyanaraman et al., 2017). In contrast, oxidative modification of PKG1-1 (but not PKG1-2) was previously suggested to result in activation of the kinase in a cGMP-independent manner (Burgoyne et al., 2007b), but we were unable to detect such an effect in vitro (fig. S9).

CAMK1 has been reported to be inhibited by glutathionylation of Cys179 in the activation segment (Kambe et al., 2010). Consistently, we found that oxidation was sufficient to inactivate two isoforms of CAMK1, CAMK1A and CAMK1D, both of which were assayed in an identical fashion (Fig. 7H-I). Inactivation is potentially as a consequence of Cys179 oxidation, and likely independent of the GST tag, which was removed proteolytically in the CAMK1A sample, but still present in CAMK1D. Consistently, DTT exposure reversed H_2_O_2_-dependent inhibition of both CAMK1 isoforms (Fig. 7H-I). The related kinase CAMK2 lacks an activation segment Cys residue, but redox-regulation of dual-Met residues has been reported to promote CAMK2 activation by stabilizing a calcium/calmodulin-independent species (Erickson et al., 2008). Consistently, CAMK2G, was robustly activated by DTT, and inhibited reversibly by H_2_O_2_ (Fig. 7K), suggesting a non Cys-activation segment mechanism of redox regulation for this kinase. In contrast, CAMK2D was completely resistant to experimental reducing and oxidizing conditions when assayed in the presence of Ca^2+^ and calmodulin (Fig. 7J), demonstrating that neither peroxide nor DTT act as general regulatory factors for recombinant kinases under these specific assay conditions. Taken together, our data confirm that redox regulation is a conserved, reversible mechanism for many Ser/Thr protein kinases, and might be much more common than previously anticipated, especially amongst CAMK and AGC kinase family members containing activation segment Cys residues.

Finally, we examined redox regulation in Tyr kinases, which in contrast to regulatory phosphorylation sites, do not contain conserved Cys residues in the activation segment (Fig 5A). EPHA3 was completely resistant to inhibition or activation by peroxide and DTT respectively (fig. S10A). ABL also lacks a T-loop Cys residue but is susceptible to alternative modes of redox-dependent regulation (Leonberg and Chai, 2007). Reversible inactivation of ABL by peroxide was readily identified (fig. S10B). We next attempted to sensitize EPHA3 to Cys-based redox regulation by incorporating Cys residues at equivalent positions in the activation segment (Fig. 5C). However, both G784C and a double mutant EPHA3 protein containing G783C and G784C substitutions remained unresponsive to peroxide or DTT (fig S10D, E). G783C ABL lacked detectable phosphotransferase activity (fig. S10C).

## DISCUSSION

Aurora A is modulated by finely-tuned regulatory pathways involving both allosteric and reversible phosphorylation-dependent mechanisms, which together control catalysis (Eyers and Maller, 2004, Eyers et al., 2003, Dodson and Bayliss, 2012a, Dodson et al., 2013). Aurora A is related to the AGC protein kinases, and several of the regulatory principles that control AGC kinase catalysis (Pearce et al., 2010, Manning et al., 2002, Wilson et al., 2018) are also conserved in Aurora kinases. Redox dependent modes of regulation involving reactive Cys residues have previously been reported in other AGC kinases, such as PKA and AKT (Humphries et al., 2002, Tan et al., 2015, Truong and Carroll, 2013, Corcoran and Cotter, 2013b). In this study, we demonstrate for the first time that Aurora A is also susceptible to reversible, oxidative-inactivation in vitro and in cells, which we show is due to a reactive cysteine (Cys 290) located in the conserved activation loop, two amino acids C-terminal to the regulatory site of T288 autophosphorylation. Careful biochemical analysis of Aurora A positioned us to characterize an evolutionarily conserved, redox-dependent signaling pathway, for other Ser/Thr kinases possessing analogous Cys residues.

### Aurora A is a redox-sensitive Ser/Thr kinase

In our initial Aurora A biochemical analyses we investigated the ability of ROS, such as H_2_O_2_, and the sulfhydryl-specific oxidant, diamide (Kosower and Kosower, 1995), to modulate the catalytic activity of Aurora A in real-time. We demonstrated potent inhibition of Aurora A-dependent phosphorylation of both peptide and protein substrates following exposure to oxidizing agents. Importantly, activity could be rescued using reducing agents such as DTT, and chemical reduction of purified Aurora A purified in the presence or absence of DTT robustly enhanced kinase activity (Fig. 1). The observed oxidation-dependent inhibition of Aurora A was not judged to be a consequence of protein destabilisation or disruption of ATP binding, but instead was a direct result of oxidative modification of Cys 290 reactive cysteine residue(s). Oxidative Cys modifications, such as protein sulfenylation and sulfenamide, have distinct chemical attributes that can be detected with chemoselective probes such as dimedone (Maller et al., 2011, Forman et al., 2017). We used an antibody with specificity towards Cys-SOHs that have been covalently derivatized with dimedone, to monitor alterations in Aurora A SOH content. These data unequivocally demonstrated direct oxidative modification of Aurora A at Cys residues after exposure to H_2_O_2_ resulting in an increase in total protein sulfenylation (Fig. 2). Interestingly, not all Cys residues are equally susceptible to oxidation in proteins, with those that have a functional role in redox-dependent signaling often possessing a low pKa value of approximately 5.0 (Brandes et al., 2009). In addition, the solvent accessibility and the structural micro-environment impact the reactivity of Cys-residues (Brandes et al., 2009, Soylu and Marino, 2016). In addition to SOH, reversible Cys-sulfenamides have also been implicated as targets of dimedone adduct formation, and we are currently undertaking detailed investigation to formally distinguish between these two reversibly oxidized thiol species (Forman et al., 2017) in Aurora A.

Direct *S*-glutathionylation of Aurora A was also detected at Cys 290 (Fig 3), which is consistent with disulfide exchange between GSSG and reactive Cys thiolates in the kinase (fig. S1). The presence of sulfenylated (or sulfenamide) Cys in Aurora A was of particular interest, since this is a reversible Cys-modification and might therefore function as a bona fide signaling mechanism in response to cellular oxidative stress, as previously described for protein Tyr phosphatases (Tonks, 2005). Our observation that C290A Aurora A was still modified by dimedone, albeit to a lesser extent than for control Aurora A (Fig. 2), provides evidence for the existence of additional redox active Cys residues, although their relevance for regulating catalytic activity appears to be very minor. Interestingly, a very recent study found that Aurora A can be covalently modified by the sulfhydryl moiety of the Cys-containing metabolite CoA under appropriate redox conditions, forming a disulfide bond with Cys 290 to inhibit Aurora A activity competitively with the binding of ATP (Tsuchiya, 2018).

To better understand the molecular basis of Aurora A redox-sensitivity we utilized Cy-preds, an automatic web algorithm for the analysis and prediction of reactive Cys residues in proteins (Soylu and Marino, 2016). Using this approach, which combines structural energetics and similarity-based considerations, we identified Cys 290 in Aurora A as a very high-probability target for oxidation. The location of this residue on the activation loop (or T-loop) of Aurora A is notable, as this region is a known regulatory region for the modulation of catalytic activity in many eukaryotic protein kinases (Beenstock et al., 2016, Pearce et al., 2010). We propose that Cys 290 in Aurora A is strategically positioned to trigger a regulatory response to ROS, or for protection from over-oxidation by covalent glutathionylation. This hypothesis has previously been explored for equivalent activation loop Cys residues including Cys 200 of PKA (Humphries et al., 2002, Humphries et al., 2007a), Cys 310 of AKT (Murata et al., 2003a), Cys174 of AMPK (Shao et al., 2014), and Cys179 of CAMK1 (Kambe et al., 2010). Structural analyis of AKT has revealed an oxidized intramolecular species, likely to be the catalytically-inactive version present upon exposure to peroxide in our studies (Fig. 7A and fig. S11). Together, our findings establish Cys 290 as a key target of oxidative modification and a dominant coordinator of redox regulation in Aurora A. Although it is unlikely that Cys 290 directly participates in substrate phosphorylation by Aurora A, maintaining a reduced sulfhydryl at this position appears to be required for enzyme activity in the absence of allosteric regulators such as TPX2, which can protect Aurora A from oxidative inhibition in vitro (Fig. 2).

### Cellular modulate of Aurora A activity by reversible oxidation

The findings from our biochemical studies are supported by cellular data. We observed inhibition of TACC3 Ser558 phosphorylation (a physiological biomarker for Aurora A activity) in cells exposed to both oxidants and inducers of oxidation (Fig. 4). In addition, inhibition of Aurora A by H_2_O_2_ could be blocked by including ROS-scavengers such as NAC and GSH in the culture medium, presumably by restoring oxidized Aurora A back to a reduced, catalytically-active state or by protection of Aurora A oxidation by a TPX2-like mechanism (Fig. 2A). Interestingly, oxidative stress has been shown to impede mitotic progression of cells via a number of different mechanisms (Burhans and Heintz, 2009, Chiu and Dawes, 2012). To ensure that changes in signaling were a direct consequence of oxidative modification of Aurora A, and not just due to cell cycle inactivation of the kinase, all of our experiments were performed using synchronized cells arrested in mitosis with nocodazole, which is itself associated with an elevation in the amount of Cys sulfenic acid present in proteins (Patterson et al., 2019). Previous observations demonstrated hyperphosphorylation of Aurora A at Thr 288 in HeLa cells under oxidative stress (Wang et al., 2017), which was proposed as a mechanism for ROS-induced mitotic arrest. Although Thr 288 autophosphorylation is considered a critical kinase activating step, a growing number of studies now suggest that this parameter is non-ideal for reporting Aurora A activity in cells (Shagisultanova et al., 2015). In this regard, it is noteworthy that phosphorylated Thr 288 can potentially be generated by non-autophosphorylation mechanisms (Walter et al., 2000b, Zhao et al., 2005). Furthermore, Aurora A regulation is a dynamic, multi-layered process involving several regulatory steps that are uncoupled from autophosphorylation, including allosteric activating complexes formed with non-catalytic protein binding partners (Willems et al., 2018). Based on our data, we propose that Aurora A-dependent phosphorylation of TACC3 at Ser 558 is an ideal biomarker for reporting Aurora A redox-regulated activity, since Aurora A is the only kinase known to target this site physiologically. In support of this finding, an increase in Aurora A autophosphorylation and CoAlation is also found in cells treated with inducers of oxidative stress (Tsuchiya, 2018). Regardless of the mechanism, complex spatio-temporal regulation means that caution should be applied when interpreting changes in cellular Aurora A catalytic outputs, especially if changes in the redox environment are induced or suspected.

The redox regulation of signaling enzymes is a rapidly emerging field. Whereas the majority of early research focused on the indirect targeting of kinases through oxidative inhibition of protein tyrosine phosphatases (Brandes et al., 2009), there is now a wealth of evidence detailing direct oxidation of Cys and Met residues in protein kinases, where diversity amongst kinase groups and subfamilies has been reported (Corcoran and Cotter, 2013b, Truong and Carroll, 2013). However, although Cys-dependent redox regulation has been described within stress-activated protein kinase modules, including thioredoxin-regulated ASK1 (Saitoh et al., 1998, Nadeau et al., 2009, Park et al., 2004), MEKK1 (Janet and Templeton, 2004), MKK6 (Diao et al., 2010), and glutathione-responsive JNK and p38α-MAPK (Wilhelm et al., 1997), no conserved mechanism has been described, to our knowledge. Moreover, none of these redox-regulated kinases contain an appropriate activation segment Cys residue (see below), in contrast to the physiological p38-MAPK downstream targets MAPKAP-K2 and MAPKAP-K3, which we conclude are rapidly inactivated by oxidation in vitro (Fig. 7).

### Evolutionary bioinformatics reveals that Cys is widespread in ePKs

A comparative evolutionary analysis of protein kinomes revealed that ∼11.5 % of all protein kinases contain an analogous Cys residue to Aurora A Cys 290 (equivalent to Cys 200 in PKA) at the ‘T-loop +2 position’ in the activation segment (Fig. 5). However, only specific members of the AGC and CAMK sub-families were enriched for this conserved Cys residue. This prompted us to search for redox-sensitivity in a selection of kinases containing a conserved T-loop Cys. Of 17 ‘T-loop +2 Cys’-containing kinases investigated, 13 were susceptible to reversible oxidative modulation. These included kinases for which redox-sensitivity had previously been noted, including AKT, AMPK, MELK and CAMK1, and, to our knowledge, novel targets of oxidative modification, including SIK1-3, PLK1, PLK4 and MAPKAP-K2/3. Intriguingly, some kinases that were predicted to be redox-sensitive based on the presence of the appropriate Cys-residue, such as Protein Kinase G (PKG) and Phosphorylase Kinase (PhK), were extremely resistant to peroxide inhibition, under the same experimental redox conditions that led to reversible modulation of other kinases. A lack of chemical reactivity in an activation segment Cys might partially explain this observation, which also confirms that in our standard assay these concentrations of redox reagents are not inducing effects through a non-specific mechanism, such as protein denaturation. The intrinsic pKa value of individual Cys residues, and their susceptibility to oxidation, is influenced by networks of interacting amino acids, solvent accessibility, protein-protein interactions and structural dynamics (Soylu and Marino, 2016, Poole, 2015). In this context, artificial incorporating of a Cys residue at the equivalent position in EPHA3 did not sensitize this Tyr kinase to oxidation (fig. S10), suggesting that the EPHA3 activation loop is an unfavourable environment to stabilize reactive Cys residues. Moreover, the relative reactivity of a Cys-residue is likely to be context-specific and vary between different allosteric activation states. In this regard, it is noteworthy that Cys 290 transitions from being exposed, to buried, in TPX2-bound Aurora A (Tsuchiya, 2018) although it is not immediately obvious to what extent this reconfiguration translates to a reversible change in Cys reactivity.

### The Cys-containing regulatory activation segment in ePKs

The conserved location of Cys residues described in this study is specific to CAMK and AGC ePKs, although well-studied groups of kinases in these families such as G-protein coupled kinases (GPRKs), PDK1, NDR/LATS kinases, MLCKs or DAPKs do not contain a Cys at the Cys 290 equivalent of Aurora A. Closer inspection of the activation segment confirms that although Cys is present at all possible positions in the activation segment in various kinomes (fig. S7A), two sites, DFG +2 (5.1%, fig. S7A) and T-loop +2 (11.5%, Fig. 5A) dominate evolutionary Cys conservation. Interestingly, both Cys residues acids are co-conserved in ∼1.4% of all ePKs in databases (Fig. 5A, bottom panel), where in the context of redox regulation, they are known to support intramolecular and/or intermolecular disulfide bonds in AKT and MELK (fig. S11). The kinase activation segment (Fig. 5A, top) contains multiple conserved residues available for post-translational phosphorylation in protein kinases (McSkimming et al., 2016, Caron et al., 2016b, Wilson et al., 2018, Cobbaut et al., 2017), and serves as a critical regulatory structure for the modulation of catalytic activity (Nolen et al., 2004). The activation loop of Aurora A itself undergoes dynamic conformational changes in response to phosphorylation and interactions with allosteric binding-partners, enabling Aurora A to transition between alternate active states (Levinson, 2018). This structural plasticity is perhaps most apparent when considering the number of distinct Aurora A conformations that have been captured in complex with small molecule inhibitors by crystallographic and NMR based techniques (Dodson et al., 2010, McIntyre et al., 2017, Pitsawong et al., 2018, Lake et al., 2018). Even in an active, phosphorylated form, the activation loop of Aurora A possesses a dynamic conformational ensemble of ‘DFG-in’ sub-states (Gilburt et al., 2017, Ruff et al., 2018). We postulate that oxidation of Cys 290, or equivalent Cys residues in the activation segment, directly influence activation loop topography to promote a switch to a less active sub-population, or are involved in the generation of higher-order molecular species such as inhibitory dimers. The observation that activation by TPX2 binding to Aurora A supersedes oxidative inhibition is consistent with this explanation, given that inactive (e.g Thr 288 dephosphorylated) Aurora A assumes an active conformation when it is bound to TPX2 (Ruff et al., 2018). Clearly however, further efforts are required to decipher the molecular and structural processes involved in oxidative regulation of protein kinases and to assess the relative contribution of different reversible Cys oxidized states (e.g sulfenic acids, sulfenyl-amides, intra-and intermolecular disulfide bonds) in modulating function. The close proximity of the most conserved Cys residues in ePKs, is known to enable a regulatory intramolecular disulfide bond to form, including Cys296 and Cys310 in the auto-inhibited conformation of AKT and Cys153 and Cys 168 of MELK (Huang et al., 2003, Murata et al., 2003a, Cao et al., 2013a). MELK also provides a plausible mechanistic explanation for the oxidative inactivation of structurally homologous kinases through dimerization (fig. S11). Although MELK kinase activity exhibits an obligate dependence on the presence of reducing agents (Wang et al., 2014), elimination of the intramolecular disulfide bond by double mutation of both Cys residues releases MELK from this regulatory requirement (Cao et al., 2013a). Interestingly, in our experiments removal of either Cys168 or Cys169 (the latter of which represents the T+2 Cys) was insufficient to abolish redox dependent activation of MELK, which potentially indicates a redundancy in the ability of both residues to form a disulfide bond with Cys153. This was substantiated with a double Cys168/169 MELK mutant, which was no longer activated by DTT. For T+2 Cys kinases that lack a complementary Cys residues with which a disulfide bond could form, the molecular mechanisms of redox based regulation are less apparent, although we cannot discount inter-or intradisulfide bond formation between other, as yet uncharacterized, Cys residues.

## CONCLUSIONS

In this paper, we describe a conserved mechanism for the oxidative modulation of protein kinase activity involving structurally homologous redox-active Cys residues located on the activation loop. Due to a central role in controlling catalytic output through phosphorylation, this region of kinases has been heavily investigated as a regulatory hot-spot. Our new findings suggest that Cys oxidation and reduction act dominantly to T-loop phosphorylation, providing an extra phosphorylation-independent layer of control over enzyme activity. The activation segment has been exploited for the generation of many phosphospecific antibodies for biological evaluation CAMK and AGC kinases. To our knowledge, the effects of Cys redox status have not been tested in the context of phosphospecificity, and in most cases, Cys residues in proteins are reduced by boiling in buffer containing a reducing agent prior to SDS-PAGE. However, our work demonstrates the vulnerability of Cys residues to redox modification in human AGC and CAM kinases, which impinges directly on catalytic output, and might also impact on the ability of antibodies to accurately monitor phosphorylation status of the activation segment.

Our study opens new investigative avenues to explore the functional relationship between physiological ROS-based signaling networks and the broader redox-regulated kinome. Mitochondrial damage and the associated elevation of ROS is implicated in a range of human diseases including ageing (Bharadwaj et al., 2015), cancer (Wallace, 2012) and neural degenerative disorders such as Parkinson’s disease (Arun et al., 2016) and a multitude of factors, including hypoxia, contribute to sustained high ROS levels in tumours (Gill et al., 2016). A key line of enquiry, therefore, will be to explore the impact of chronic oxidative stress and increased sulfenylated-protein populations on both normal and pathological Ser/Thr kinase function. The extent to which oxidation of kinases may influence the therapeutic efficacy of inhibitor compounds in a cellular context is also of potential interest, especially for kinases targeted by Cys-based covalent mechanisms. Indeed, different Aurora A inhibitors target distinct conformational species, which can be broadly separated into compounds with preferences for the DFG-in or DFG-out states (Lake et al., 2018, Pitsawong et al., 2018, Ruff et al., 2018). The ability of oxidative modifications in the activation segment to alter this ‘DFG equilibrium’ may also have implications for the selectivity of inhibitors in cells, where the propensity of redox-active Cys residues in Ser/Thr kinases to undergo sulfenylation could be exploited for the rationale design new classes of covalent inhibitors. This strategy has been adapted to great success to generate clinical compounds with potency and selectivity towards tyrosine kinases, such as afatinib and neratinib, which target Cys797 of the redox-regulated tyrosine kinase EGFR for the treatment of non-small cell lung cancer (Singh and Jadhav, 2018). Finally, a deeper mechanistic understanding of the dynamics of Ser/Thr kinase redox regulation, may reveal Cys residues that are differentially exposed in active and inactive kinase conformations and potentially lead to a diverse and versatile reservoir of specific drug targets (Leproult et al., 2011).

## MATERIALS AND METHODS

Commercial recombinant protein kinase fusion proteins were obtained from MRC PPUU reagents (University of Dundee), and were purified from Sf21 cells or *E. coli.* Full details are provided in Table 2. GST and 6His-tagged kinases were purified using standard procedures, prior to storage in 1 mM DTT at −80°C. All kinases were assayed using standard enzyme preparations (Bain et al., 2007). Bacterially-expressed GST-MAPKAP-K2 and GST-MAPKAP-K2 were activated in vitro by incubation with catalytically-active p38a, which was subsequently removed by re-purification prior to assay. General biochemicals and all redox reagents, including glutaredoxin (GRX), were purchased from Sigma-Aldrich.

### Protein kinase assays

Kinase assays were performed using non-radioactive real-time mobility shift-based microfluidic assays, as described previously (Byrne et al., 2016, Caron et al., 2016a, McSkimming et al., 2016, Mohanty et al., 2016) in the presence of 2 μM of the appropriate fluorescent-tagged peptide substrate (Table 2) and 1 mM ATP. Pressure and voltage settings were adjusted manually to afford optimal separation of phosphorylated and non-phosphorylated peptides. All assays were performed in 50 mM HEPES (pH 7.4), 0.015% (v/v) Brij-35, and 5 mM MgCl_2_, and the real-time or endpoint degree of peptide phosphorylation was calculated by differentiating the ratio of the phosphopeptide:peptide present in the reaction. Kinase activity in the presence of different redox reagents was quantified by monitoring the generation of phosphopeptide during the assay, relative to controls. Data were normalised with respect to control assays, with phosphate incorporation into the peptide limited to ~20% to prevent depletion of ATP and to ensure assay linearity. ATP K_M_ and the concentration of a compound that caused 50% inhibition (IC50) values were determined by nonlinear regression analysis using Graphpad Prism software. Where specified, kinase assays employing Aurora A were supplemented with 100 nM GST-TPX2 or GST alone. Assays for CAMK kinases included 5 mM CaCl_2_ and 2 μM Calmodulin as standard. Where appropriate, PKG1 assays were performed in the presence of 1 mM cGMP. Recovery of Aurora A activity from oxidative inhibition was assessed by monitoring substrate phosphorylation in the presence of peroxide in real time, followed by subsequent introduction of DTT or GSH. To standardize this assay for all kinases, enzymes were pre-incubated in the presence or absence of 5 mM H_2_O_2_ on ice for 30 mins and then substrate phosphorylation was initiated with the addition of 1 mM ATP and the appropriate substrate peptide in the presence (where indicated) of 10 mM DTT. Aurora A kinase assays were also developed with recombinant GST-TACC3 as a substrate. TACC3 Ser 558 phosphorylation was detected by immunoblotting with a pSer 558 TACC3 antibody, as previously described (Tyler et al., 2007). Aurora A autophosphorylation was detected using a Thr 288 phosphospecific antibody (Tyler et al., 2007).

### Recombinant Protein purification

For enzyme and DSF assays, murine or human Aurora A, MELK (1-340), PLK1 (1-364), PLK4 (1-369), full-length PKA, EPHA3 (kinase domain and juxtamembrane region), ABL (46-515), TACC3 and TPX2(1-43) were produced in BL21 (DE3) pLysS *E. coli* cells (Novagen) with expression induced with 0.5 mM IPTG for 18 h at 18°C and purified as N-terminal His6-tag or N-terminal His6-GST tag fusion proteins by affinity chromatography and size exclusion chromatography using a HiLoad 16/600 Superdex 200 column (GE Healthcare) equilibrated in 50 mM Tris/HCl, pH 7.4, 100 mM NaCl and 10 % (v/v) glycerol. Where appropriate, recombinant protein was purified in the presence of 1 mM DTT. Cys290Ala Aurora A and equivalent Cys-Ala mutants of other kinases were generated using standard mutagenic procedures, and purified as described above (Byrne et al., 2016). To generate a phosphorylation-depleted kinase, Aurora A was co-expressed with lambda phosphatase in BL21 (DE3) pLysS *E. coli* cells prior to purification.

### Differential Scanning Fluorimetry

Thermal-shift assays were performed with a StepOnePlus Real-Time PCR machine (Life Technologies) using Sypro-Orange dye (Invitrogen) and thermal ramping (0.3 °C in step intervals between 25 and 94°C). All proteins were diluted to a final concentration of 5 μM in 50 mM Tris/HCl, pH 7.4 and 100 mM NaCl in the presence or absence of the indicated concentrations of ATP, H_2_O_2_, DTT, GSH or MLN8237 (final DMSO concentration no higher than 4 % v/v) (Byrne et al., 2018) and were assayed as described previously (Foulkes et al., 2018). Normalized data were processed using the Boltzmann equation to generate sigmoidal denaturation curves, and average T_m_/ΔT_m_ values calculated as previously described (Murphy et al., 2014) using GraphPad Prism software.

### Human cell culture and treatment of cells with redox-active agents

HeLa cells were cultured in Dulbecco’s Modified Eagle Medium (DMEM) (Lonza) supplemented with 10% foetal bovine serum (FBS) (Hyclone), 50 U/ml penicillin and 0.25 μg/ml streptomycin (Lonza) and maintained at 37 °C in 5 % CO_2_ humidified atmosphere. To examine the effects of oxidative stress on Aurora A kinase activity, cells were arrested in mitosis with 100 nM nocodazole for 16h, then treated with a range of concentrations of H_2_O_2_, menadione, or diamide for 30-60 min. To stimulate chronic oxidative stress, arrested HeLa cells were collected, washed in PBS, and fresh culture medium containing glucose oxidase at a non-toxic concentration (2 U/ml) and 100 nM nocodazole was added. Subsequently, cells were collected and harvested periodically over a 2 h time period. To investigate reversible inactivation of AurA by peroxide, arrested cells were incubated for 10 mins with 10 mM H_2_O_2_, peroxide-containing medium was removed and replaced with fresh culture medium containing 10 mM DTT, NAC or GSH or buffer control. In all assays cells were subsequently washed in PBS and harvested in bromophenol blue-free SDS sample buffer supplemented with 1% Triton X-100, protease inhibitor cocktail tablet and a phosphatase inhibitor tablet (Roche) and sonicated briefly prior to immunoblotting.

### Cell lysis, immunoprecipitation and Western blot analysis

Total cell lysates were centrifuged at 20817x g for 20 min at 4°C and the supernatant was preserved for further analysis. Samples were initially diluted 50-fold and protein concentration was measured using the Coomassie Plus staining reagent (Bradford) Assay Kit (Thermo Scientific).

### Detection of sulfenylated and glutathionylated proteins by immunoblotting

Recombinant Aurora A was incubated with 50 mM Tris/HCl, pH 7.4, and 100 mM NaCl in the presence of different concentrations of H_2_O_2_ or 10 mM DTT for 10 min. Cysteine sulfenic acid was detected by SDS-PAGE and immunoblotting after adduct formation with 1 mM dimedone for 20 mins at RT. Dimedone stocks were prepared in DMSO with a final assay DMSO concentration no higher than 4 % (v/v). To detect glutathionylation of proteins, proteins were incubated with 10 mM GSSG or GSH for 30 mins and glutathione-protein complexes were detected by immunoblotting after non-reducing SDS-PAGE.

### Identification, alignment and visualization of protein kinase-related sequences

The MAPGAPS procedure (Neuwald, 2009) was employed alongside a variety of curated eukaryotic protein kinases profiles (Kannan et al., 2007, Finn et al., 2010, McSkimming et al., 2016, Talevich et al., 2011) to identify and align eukaryotic protein kinase-related sequences from the non-redundant (NR) sequence database and UniProt reference proteome (consortium, 2015) databases (Release 2018_09). Sequences with a Cys residue at the Aurora A Cys290 equivalent position were retrieved and used for further taxonomic analysis. Taxonomic information was based on NCBI Taxonomy database (Sayers et al., 2009). Weblogo’s (Schneider and Stephens, 1990) and were generated using Weblogo Version 2.8. Amino acids were coloured based on their chemical properties. Polar amino acids (G,S,T,Y,C,Q,N) are colored green, basic (K,R,H) blue, acidic (D,E) red, and hydrophobic (A,V,L,I,P,W,M) black.

## SUPPLEMENTARY MATERIALS

**Supplementary Figure 1.**
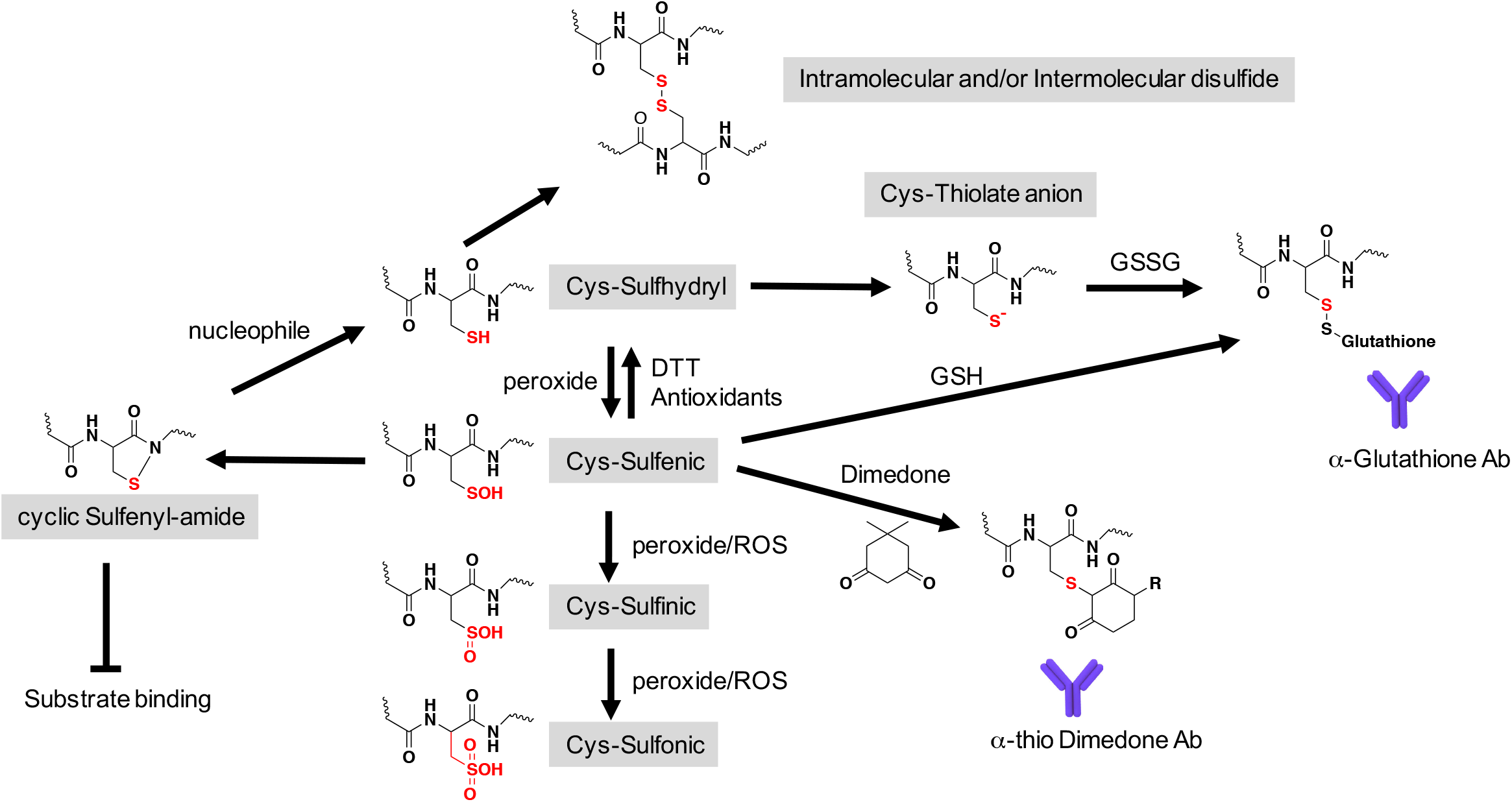
Common chemical and enzymatic routes for Cys redox modifications in proteins, and their detection with immunological reagents. A redox flow-chart is presented.

**Supplementary Figure 2.**
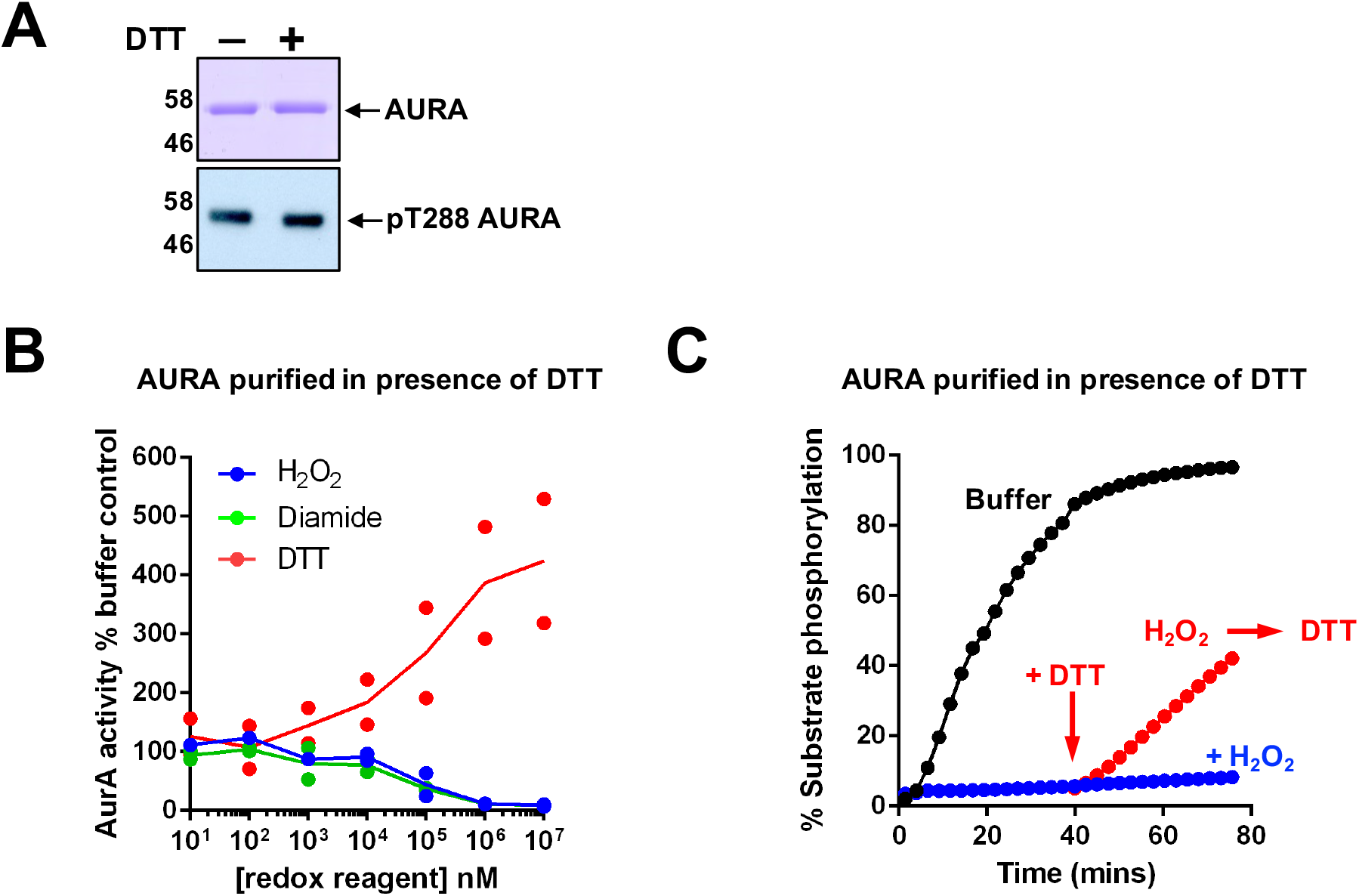
Analysis of Aurora A purified in the presence of DTT. **(A)** Recombinant Aurora A (0.5 μg each) purified in the presence or absence of DTT and resolved by SDS-PAGE and visualised by Coomassibe blue staining (top panel). Phosphorylation (pThr 288) of each protein (20 ng) was also analysed by immunoblotting (bottom panel). **(B)** Dose response curves for the reducing reagent DTT (red) and the oxidizing agents H_2_O_2_ (blue) and diamide (green) with 6 nM recombinant Aurora A purified in the presence of DTT. Aurora A activity was assessed by monitoring phosphorylation of a fluorescent peptide substrate using 1 mM ATP, and normalized to control experiments containing buffer alone after 30 min assay time. **(C)** Oxidative-inhibition of Aurora A purified in the presence of DTT is reversible. Assay conditions are as described in Fig. 1 (C).

**Supplementary Figure 3.**
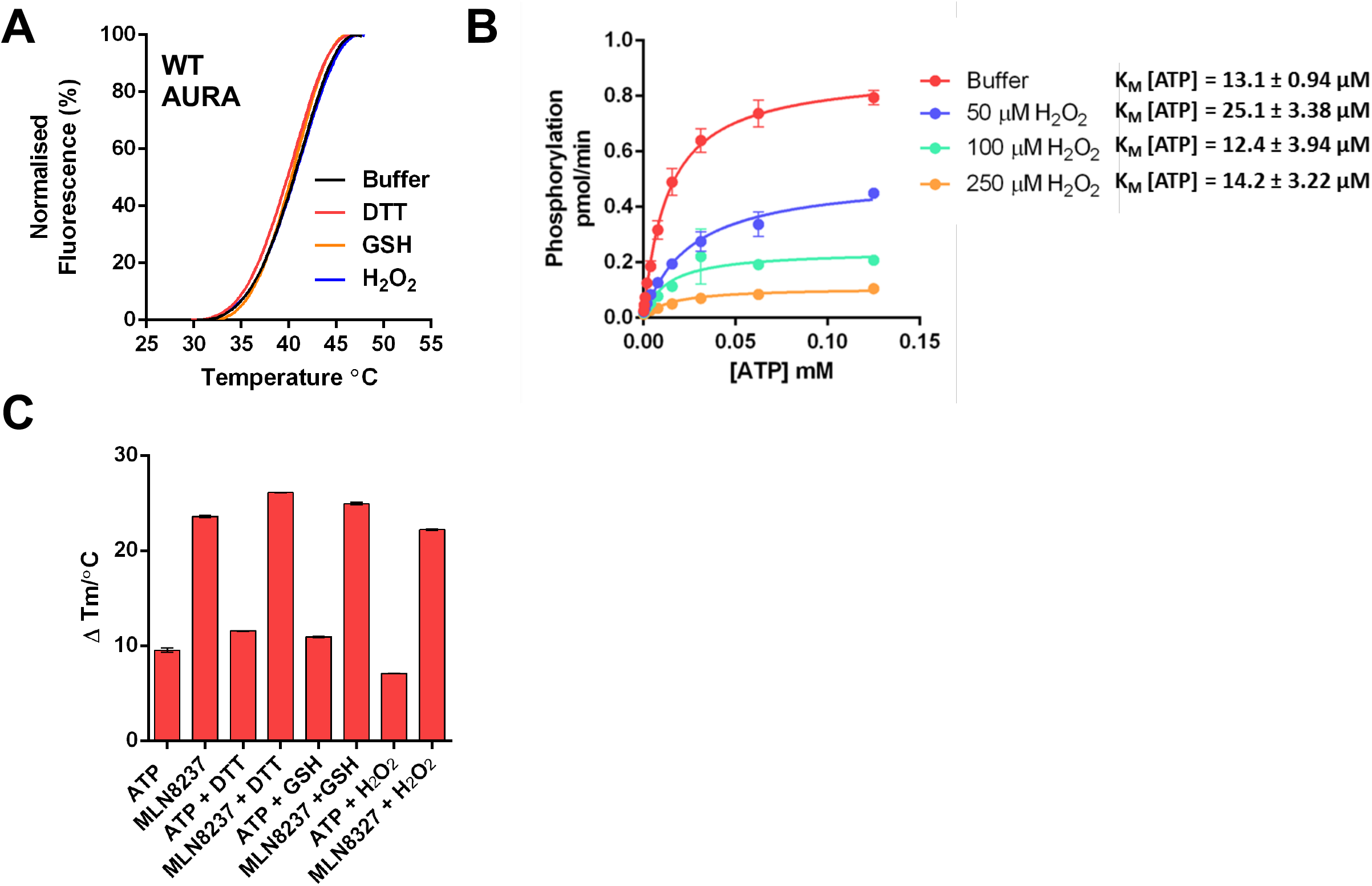
Biochemical analysis of Aurora A oxidation. **(A)** Aurora A stability is unaffected by redox state. Thermal denaturation profiles of recombinant Aurora A (5 μM) in the presence of 1 mM of the indicated redox-reagent. Representative unfolding profiles of two independent experiments shown. **(B)** K_M[ATP]_ determination for Aurora A (12.5 nM) in the presence of H_2_O_2_. Kinetic analysis of Aurora A-catalysed peptide phosphorylation (pmol phosphate/min) with increasing concentrations of ATP was performed with the indicated concentrations of H_2_O_2_. K_M[ATP]_ values (± standard deviation) were calculated from two independent experiments using GraphPad Prism software. **(C)** Analysis of thermal shifts induced by 1 mM ATP and 100 μM MLN8237 in the presence of 1 mM of the indicated redox-reagent. Means of two independent experiments shown.

**Supplementary Figure 4.**
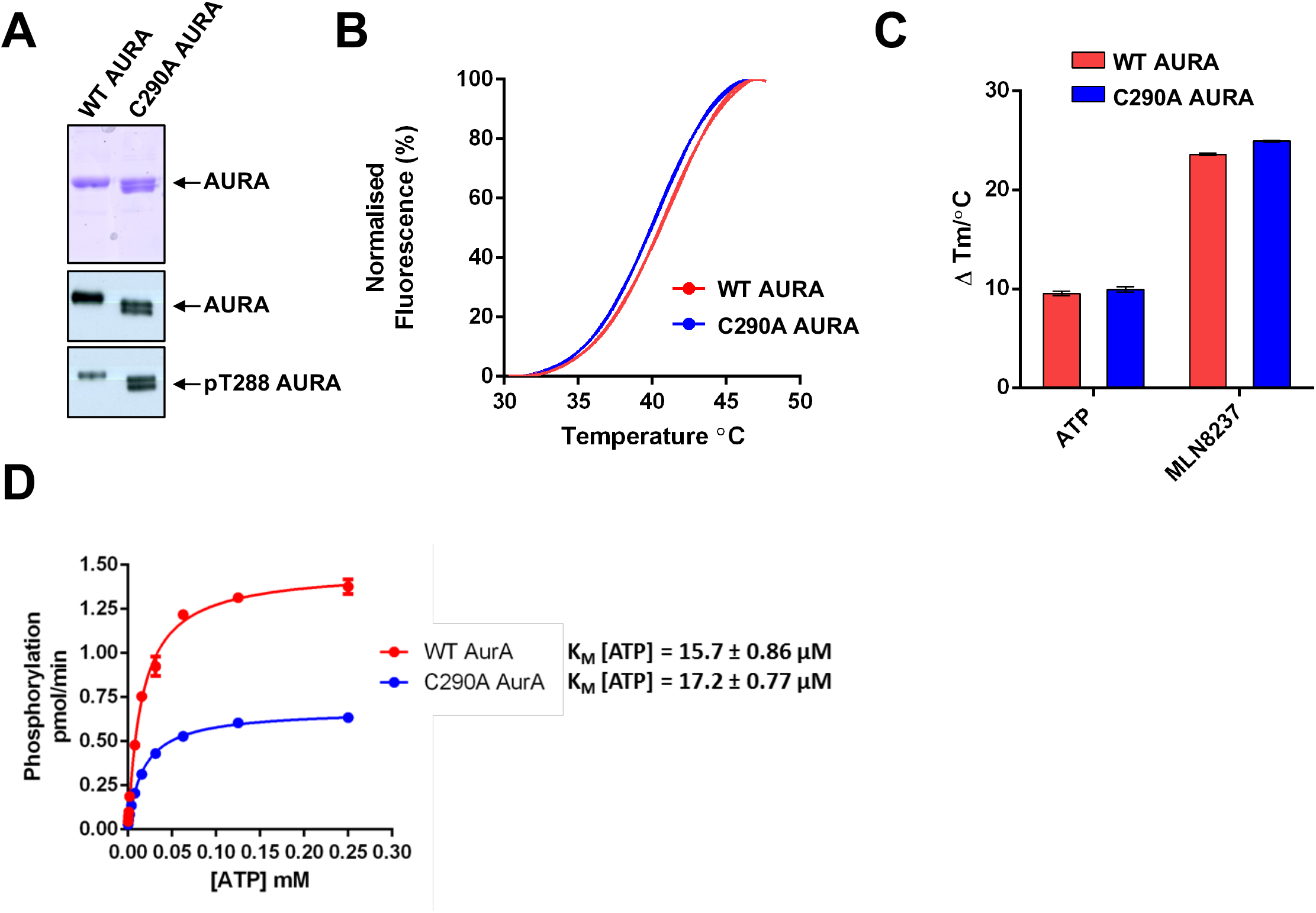
Analysis of Aurora A and a redox-resistant C290A mutant. **(A)** Purified WT and C290A Aurora A (0.5 μg each) resolved by SDS-PAGE and visualised by Coomassie blue staining (top panel) and immunoblot (50 ng protein) using antibodies with specificity for Aurora A (middle panel) and site specific autophosphorylation at Thr 288 (bottom panel). **(B)** Thermal denaturation profiles of recombinant Aurora A proteins 5 μM. Representative unfolding profiles from two independent experiments are shown. **(C)** Thermal shifts induced by 1 mM ATP and 100 μM MLN8237. **(D)** Calculation of K_M[ATP]_ values for WT (red) and C290A Aurora A (blue). ATP concentrations were varied in the presence of a fixed amount (6 nM) of Aurora A protein. The rate of peptide substrate phosphorylation (pmol phosphate/min) was calculated from two independent experiments using GraphPad Prism software.

**Supplementary Figure 5.**
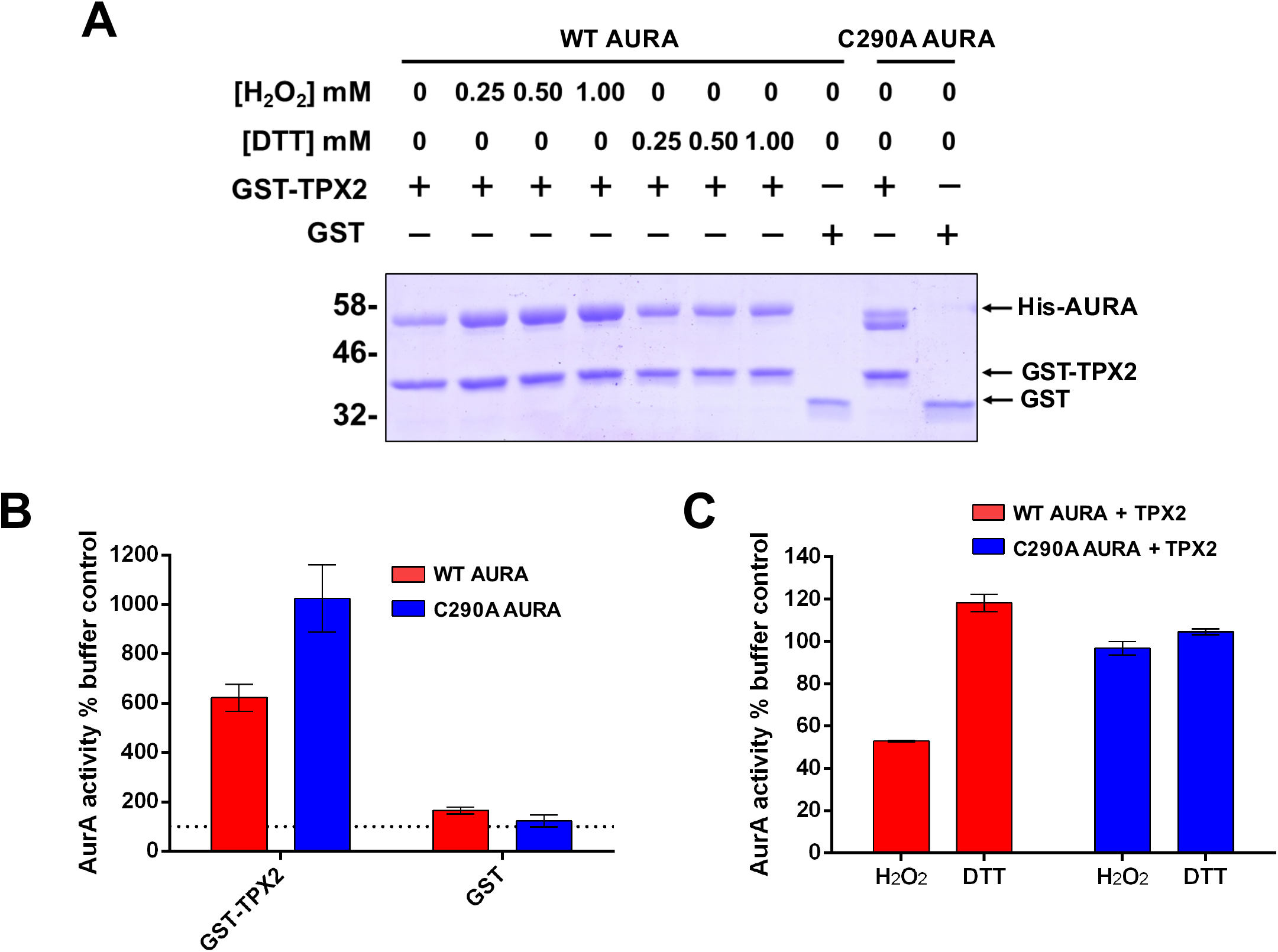
WT and C290A Aurora A both bind to TPX2. **(A)** Interaction of WT and C290A His-Aurora A with GST-TPX2 under oxidizing and reducing conditions. 5 μM WT Aurora A was incubated with the indicated concentrations of H_2_O_2_ or DTT for 5 mins at 20 °C and pulled down with GST-TPX2 glutathione beads (~1 μg GST-TPX2 per 10 μl bead volume). Protein was eluted with 10 mM GSH. Pull down assays with C290A Aurora A also shown. Control experiments used GST as a bait protein. **(B)** The activity of WT (red) and C290A (blue) Aurora A (6 nM, with 1 mM ATP) towards a peptide substrate was examined in the presence or absence of TPX2. GST at an equimolar concentration to TPX2 included as a control. **(C)** Redox regulation of TPX2-bound WT and C290A Aurora A. WT (red) and C290A (blue) Aurora A proteins were incubated with TPX2 and then assayed in the presence of 1 mM DTT or 10 mM H_2_O_2_. Activity was calculated relative to a control containing buffer only after 5 mins assay time. In all assays described here, Aurora A proteins were incubated with TPX2 for 10 mins at 20 °C before the addition of redox reagent or ATP. The concentration of TPX2 in the final reaction mixture was fixed at 100 nM.

**Supplementary Figure 6.**
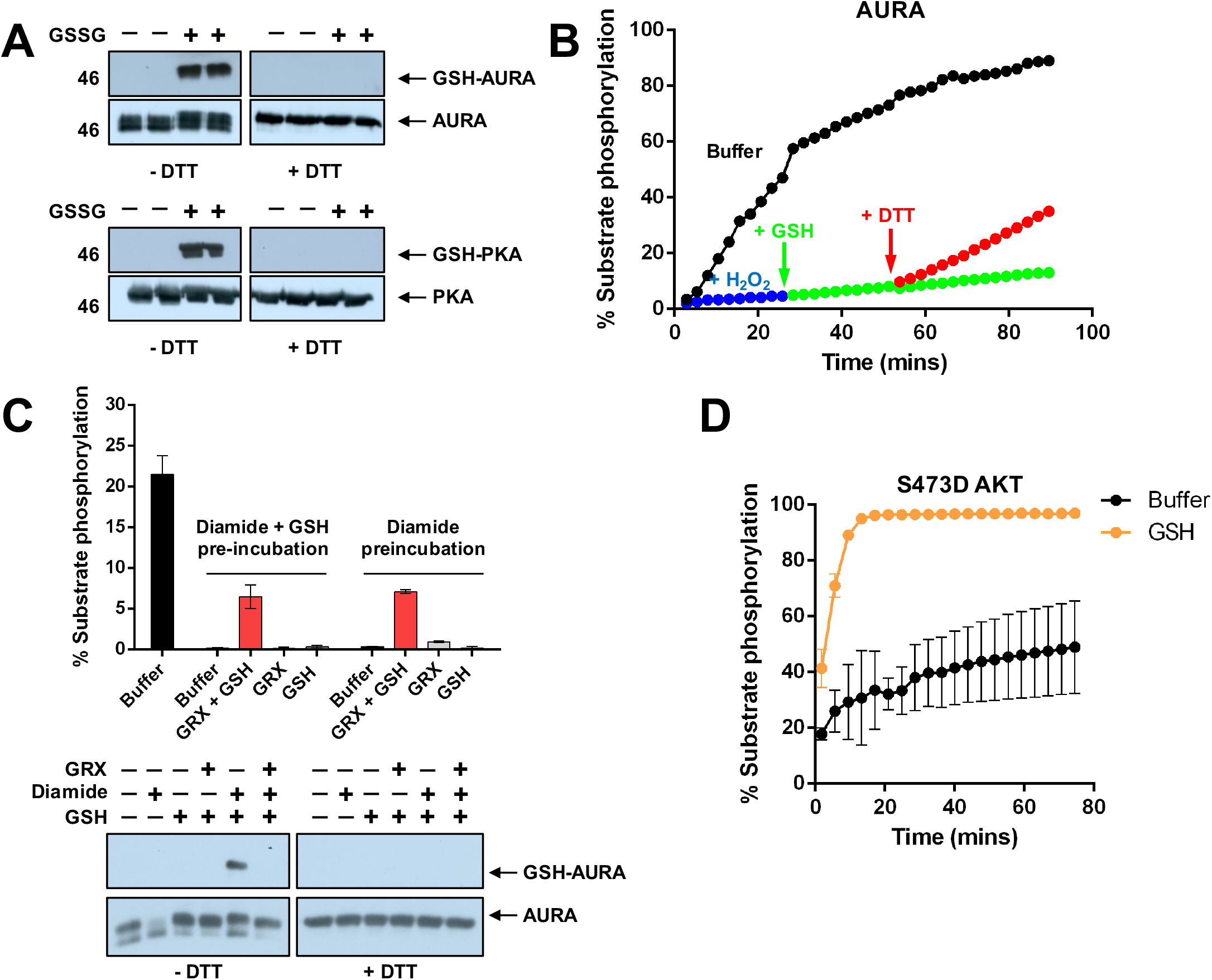
Reversible glutathionylation of Aurora A and AKT. **(A)** Immunoblot demonstrating reversible glutathionylation of Aurora A. 1 μg of recombinant Aurora A protein was incubated in the presence or absence of 10 mM GSSG for 30 mins at 20°C. SDS-PAGE was performed under reducing and non-reducing conditions. Western blots were probed with an antibody with specificity towards glutathione-conjugated proteins. Equal loading of protein was confirmed using an antibody for Aurora A. PKA (1 μg) was also included as a positive control. **(B)** Inhibition of Aurora A by H_2_O_2_ is relieved by DTT but not GSH. Aurora A (12.5 nM) activity was assayed in the presence or absence of 1 mM H_2_O_2_ for 25 mins and the reactions were then supplemented (where indicated) with 2 mM GSH, followed by 2 mM DTT after 50 mins. Aurora A-dependent phosphorylation of the fluorescent peptide substrate was initiated by the addition of 1 mM ATP. **(C)** Aurora A (40 μM) was incubated in the presence or absence of 125 μM diamide or diamide with 100 μM GSH for 20 mins at 20 °C. Following this, reactions were supplemented with buffer control, 500 μM GSH and 0.16 mgml glutaredoxin-1 (GRX), or GSH and GRX in isolation for a further 20 mins. Reactions were then sampled for western blotting analysis (upper panel) or assayed using our in vitro kinase assay (final Aurora A concentration was 200 nM, bottom panel). Data shown is mean and SD of two independent experiments. (D) AKT is potently activated by GSH. Activity of PDK1 phosphorylated S473D AKT (7 nM) was measured in the presence or absence of 2 mM GSH and 1 mM ATP.

**Supplementary Figure 7.**
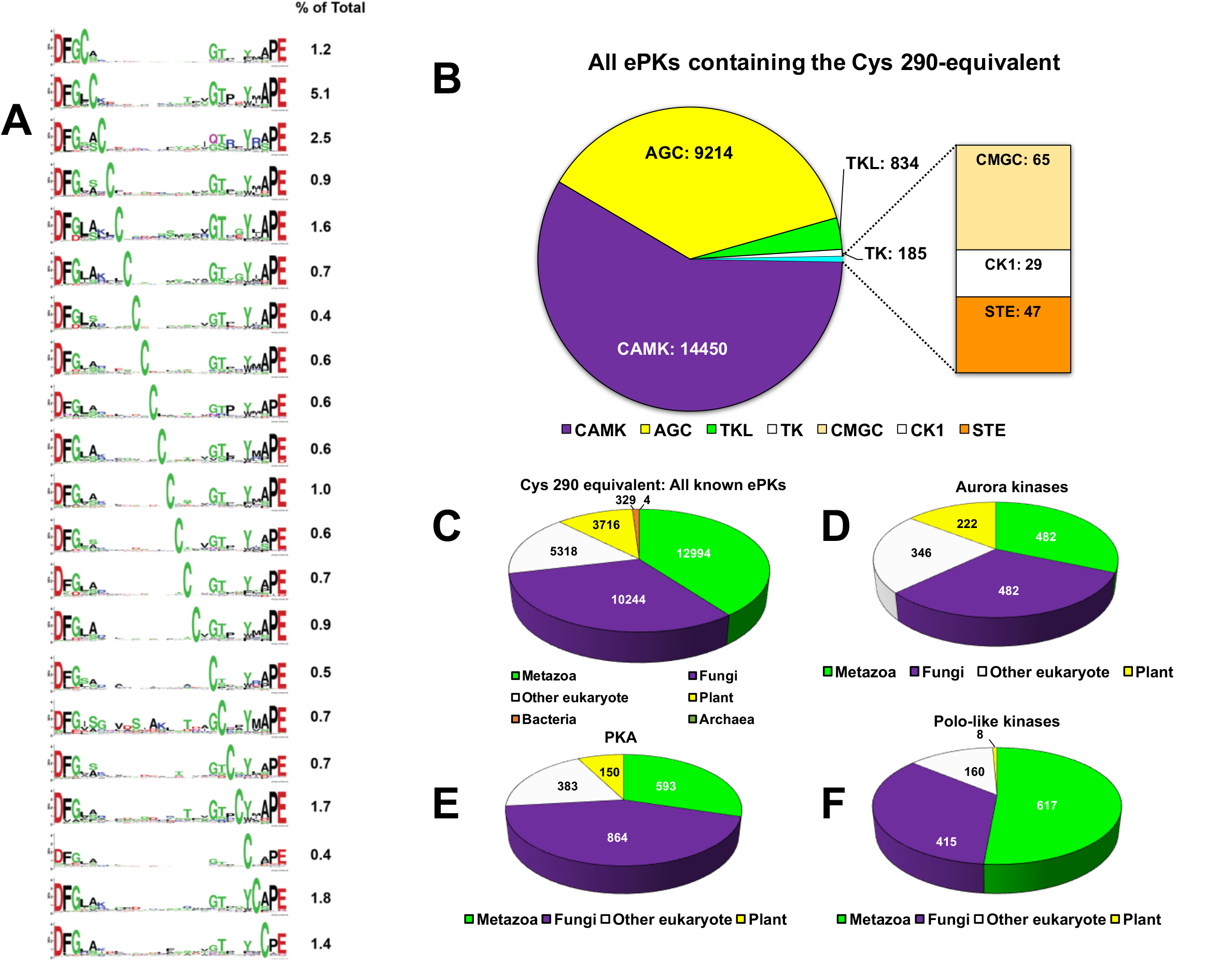
Taxonomic analysis of conserved Cys residues within the activation segment of ePKs. **(A)** The distribution of Cys residues, and co-varying amino acids at all positions within the activation segment, located between the DFG and APE residues, are displayed using WebLogos. The percentage of ePKs containing the indicated Cys residues is indicated on the right. **(B)** Kinome group distribution of all ePKs containing a Cys 290 equivalent. The total number of kinases identified within each group is indicated. **(C)** Taxonomic distribution of T loop +2 Cys-containing sequences in all ePKs **(D)** Aurora kinases, **(E)** PKA and **(F)** PLK subfamilies. Taxonomic groups are coloured as indicated and the total number of sequences identififed in each category are indicated within the pie chart.

**Supplementary Figure 8.**
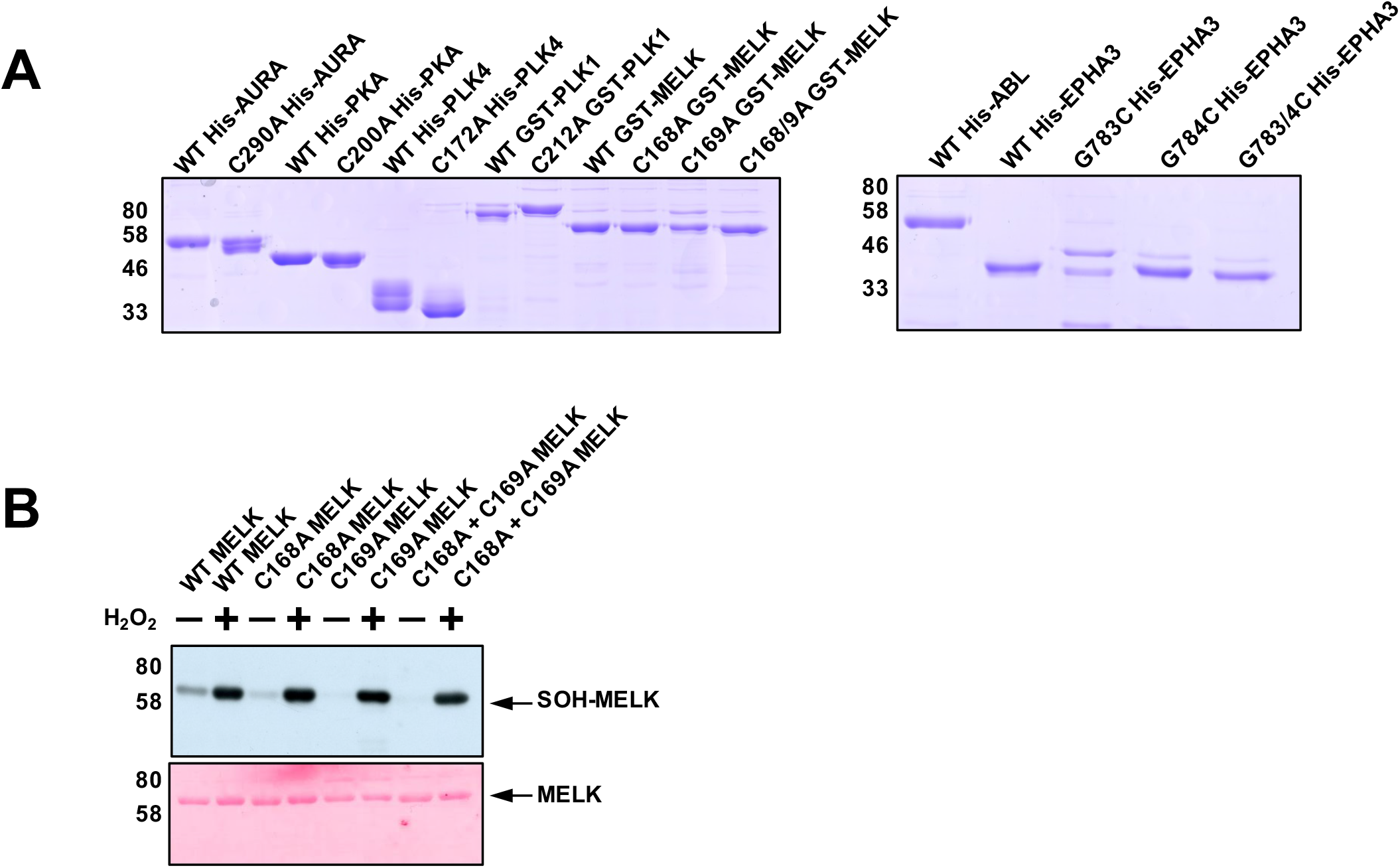
Biochemical analysis of Cys-containing protein kinases. **(A)** Purified WT and Cys-Ala mutants kinases (0.5 μg each) resolved by SDS-PAGE and visualised by Coomassibe blue staining. **(B)** Detection of reactive cysteine oxidation in WT and Cys-Ala recombinant MELK proteins with an antibody with specificity towards cysteine sulfenic acids that have been derivatised by dimedone (SOH-MELK). GST-MELK (0.5 μg) was incubated in the presence of absence of 1 mM H_2_O_2_ for 10 min and then exposed to 1 mM dimedone for a further 20 mins (all incubations performed at 20°C). Total GST-MELK loading was visualised using ponceau staining (bottom panel).

**Supplementary Figure 9.**
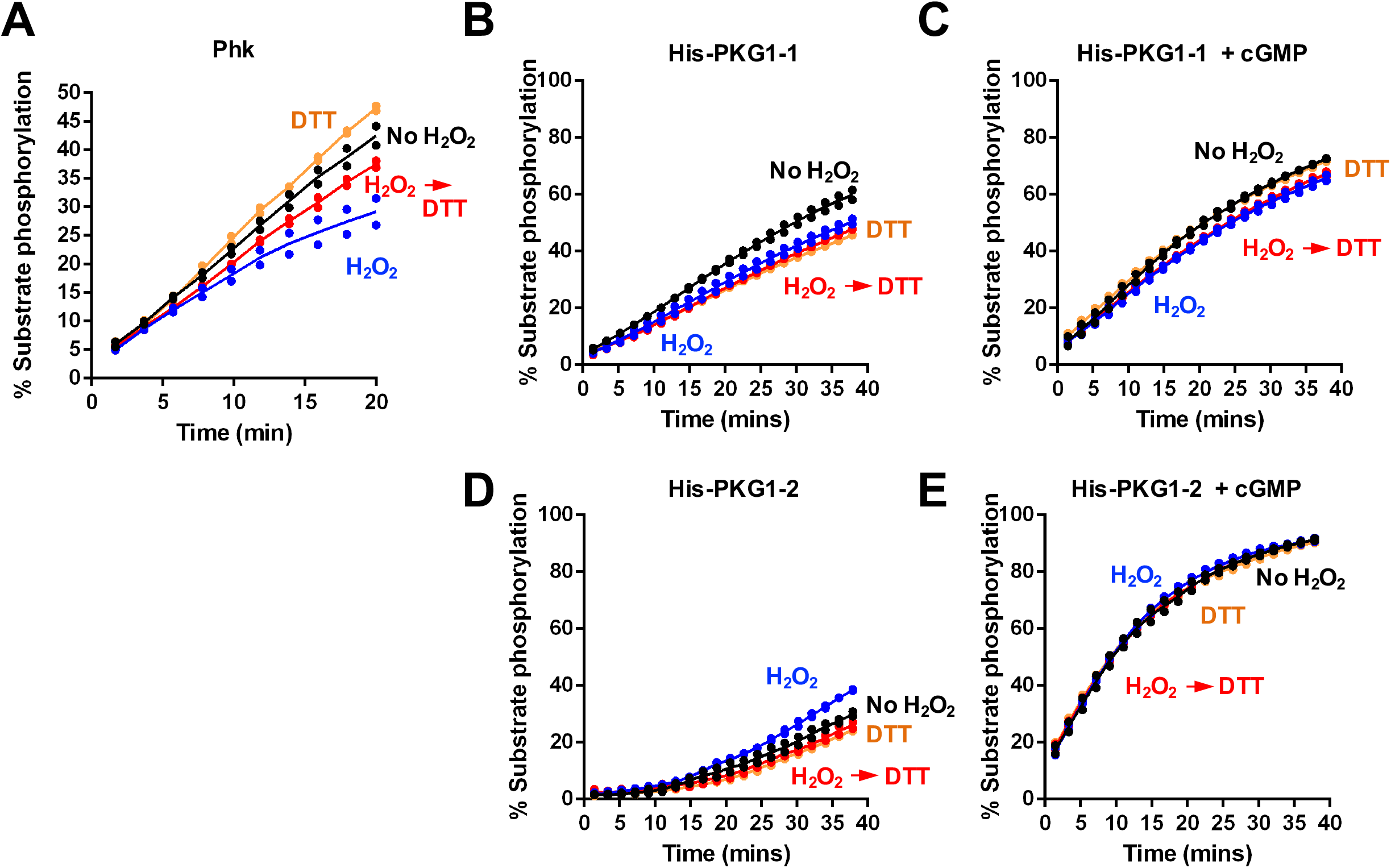
Kinases possessing a T-loop +2 Cys residue that are insensitive to redox-dependent regulation. A cohort of kinases with suspected redox sensitivity based on the presence of an analogous activation loop-located Cys residue were probed for reversible oxidation-dependent inhibition using real time phosphorylation of kinase specific peptide substrates (see Table 2). Assay conditions were as for Fig. 7 and the following concentrations of kinases were used: **(A)** 4 nM PHK, **(B, C)** 25 nM His-PKG1-1, and **(D,E)** 80 nM His-PKG1-2. Assays **(C)** and **(E)** were supplemented with 1 mM cGMP.

**Supplementary Figure 10.**
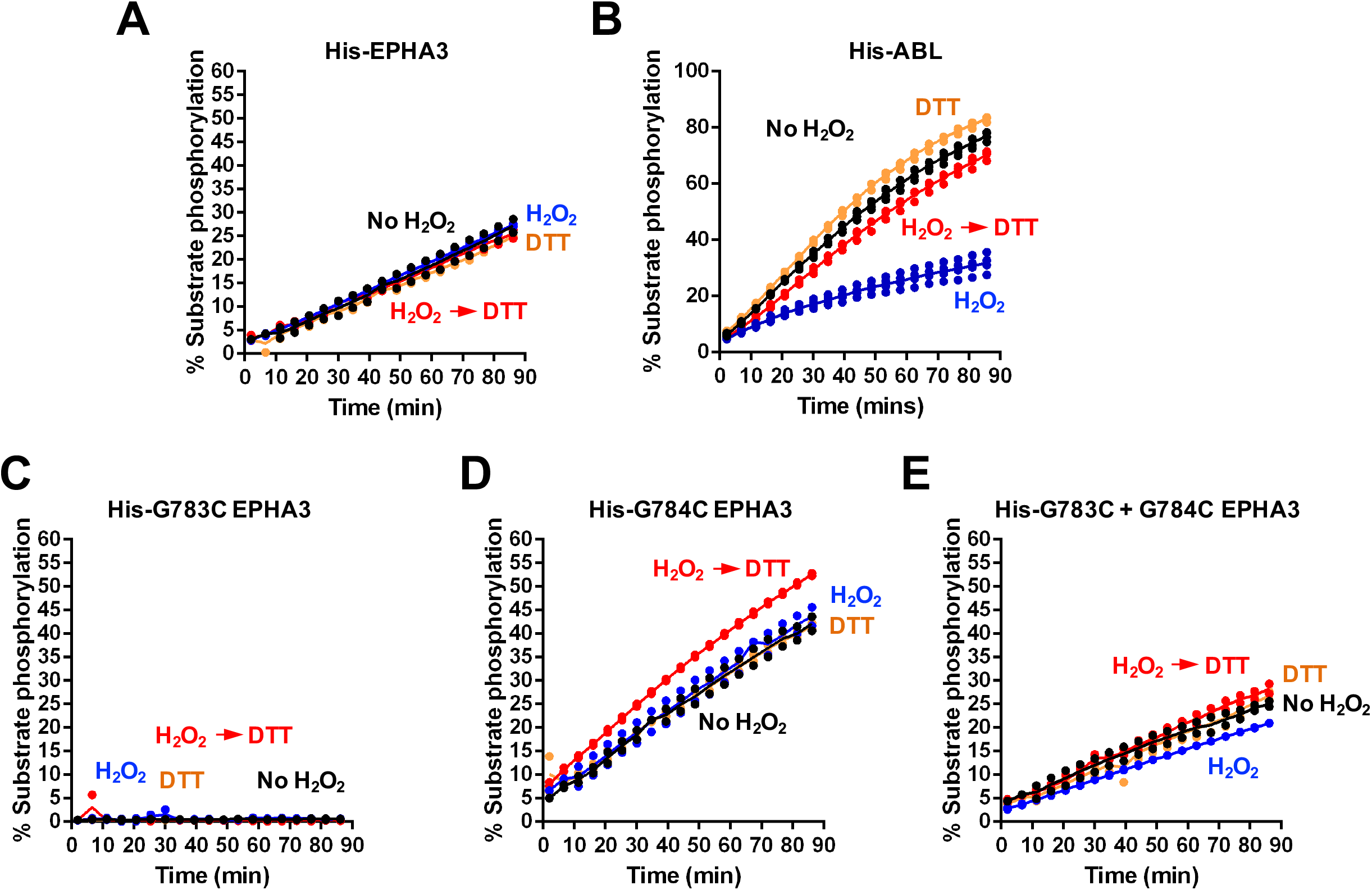
Redox regulation of Tyr kinases. Purified ABL and EPHA3 tyrosine kinases were assayed using the specified redox-dependent conditions described in Fig. 7 and Table 2. Final ABL or EPHA3 concentrations in the assay were: **(A)** 30 nM WT His-EPHA3, **(B)** 5 nM ABL and 30 nM of **(C)** G783C, **(D)** G784C or (E) G783C/G784C His-EPHA3

**Supplementary Figure 11.**
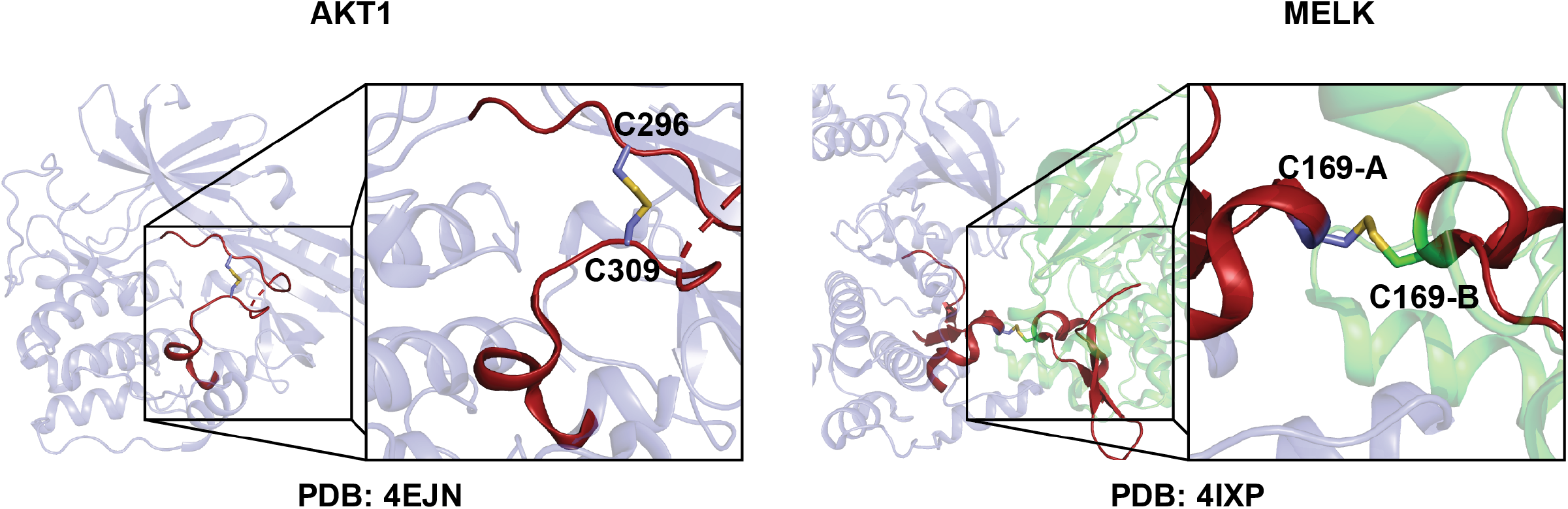
Two potential mechanisms for the regulation of Cys oxidation. **(A)** Formation of an intramolecular disulfide in AKT between Cys residues in the activation segment. **(B)** Intermolecular disulfide between exchanged activation segments of MELK

## ACKNOWLEDGEMENTS

We thank Dr James Hastie and Dr Hilary McLauchlan for help with reagents. The authors also thank Sam Evans for outstanding technical support and media preparation.

## AUTHOR CONTRIBUTIONS

PAE obtained funding and designed experiments alongside DPB, who performed all biochemical and cellular experiments. SS and NK performed bioinformatics analysis, and DPB and PAE wrote the paper, with contributions from all the authors, who together approved the final version prior to submission.

## DATA AND MATERIALS AVAILABILITY

All data needed to evaluate conclusions made are available in the main or supplementary sections of the paper.

